# Parkinson’s disease LRRK2 mutations dysregulate iron homeostasis and promote oxidative stress and ferroptosis in human neurons and astrocytes

**DOI:** 10.1101/2025.09.26.678370

**Authors:** Adamantios Mamais, Richard D. Batchelor, Aravindraja Chairmandurai, Thomas B. Ladd, Austin J. Shute, Nitya Subrahmanian, Nunziata Maio, Christopher D. Vulpe, Matthew J. LaVoie

## Abstract

**Background:** Iron accumulation is a hallmark of sporadic and familial Parkinson’s disease (PD) and correlates with clinical motor symptom severity. The biochemical mechanisms driving iron dyshomeostasis in PD brain and whether these are early or late events in the neurodegenerative process remain unknown. Nigral iron levels in LRRK2-PD patients have been reported to be even higher than in idiopathic PD, and greater in non-manifesting LRRK2 carriers than controls, suggesting that iron accumulation precedes clinical onset in LRRK2-associated PD. However, the cells affected and mechanisms governing iron dyshomeostasis in PD remain unclear.

**Methods:** Here, we investigated multiple independent measures of iron homeostasis in iPSCs, iPSC-derived cortical and dopaminergic neurons, and astrocytes from control and PD patient-derived iPSCs and an isogenic iPSC panel of three independent pathogenic PD mutations. High-content and super-resolution microscopy of iron-specific probes and ICP-MS were used to determine iron content and distribution across different cell types and LRRK2 genotypes. The upstream effectors and downstream consequences of iron dyshomeostasis on ferroptosis signaling were also examined.

**Results:** We found that heterozygous LRRK2 mutations dysregulate cellular iron levels across iPSCs, neurons and astrocytes, in a kinase-dependent manner. Lysosomal ferrous iron was specifically and consistently elevated across iPSCs, iPSC-derived cortical and dopaminergic neurons, and astrocytes carrying LRRK2 mutations and rescued by treatment with the selective LRRK2 inhibitor, MLi-2. Importantly, we show that lipid peroxidation and ROS levels are elevated in isogenic LRRK2 mutant neurons, while iron chelation or MLi-2 reduced LRRK2-dependent ROS damage. LRRK2 regulates the function of over a dozen Rab GTPase proteins through direct phosphorylation, and our prior work revealed a unique correlation with Rab8a trafficking and iron. Here, we report that CRISPR/Cas9 knockout of Rab8a recapitulates LRRK2-driven effects on intracellular iron in different cell types and that exogenous Rab8a normalizes lysosomal iron levels in R1441C LRRK2 iPSCs.

**Conclusions:** Together, our findings demonstrate that LRRK2 mutations disrupt iron homeostasis across multiple cell types, including dopaminergic neurons, and establish a discrete biochemical pathway linking LRRK2 and vulnerability to ferroptosis signaling.

## Background

Iron accumulation in the substantia nigra is a well-recognized pathological hallmark of Parkinson’s disease (PD), observed in both sporadic and familial forms of the disorder and in other neurodegenerative conditions (1–7). Iron overload correlates with neuromelanin loss in post-mortem PD brains (8) and is detectable *in vivo* through advanced MRI techniques, such as quantitative susceptibility mapping (QSM), where iron deposition in the substantia nigra pars compacta (SNc) has been correlated with motor symptom severity (9). Notably, iron-sensitive imaging aids in distinguishing PD from other parkinsonian syndromes such as multiple system atrophy (MSA) and progressive supranuclear palsy (PSP) (10–14). Despite decades of evidence linking iron dyshomeostasis to PD, the molecular underpinnings that drive brain iron accumulation and neuronal vulnerability and whether these are early or late features of disease remain incompletely understood.

Iron’s potential to promote oxidative stress through Fenton chemistry and the generation of reactive oxygen species (ROS) make it a likely contributor to dopaminergic (DA) neuron degeneration (15,16). Ferroptosis, a regulated form of iron-dependent cell death, has emerged as a key mechanism linking iron overload to neuronal loss (17–19) and has been widely implicated in PD (20–24), but how genetic factors for PD intersect with ferroptotic pathways is not fully defined. Mutations in leucine-rich repeat kinase 2 (LRRK2) are the most common cause of familial PD (25) and have been implicated in vesicular trafficking defects and lysosomal dysfunction (26–32), processes that could plausibly disrupt iron handling. Indeed, clinical imaging studies reveal that individuals carrying pathogenic LRRK2 mutations, whether symptomatic or asymptomatic, exhibit elevated nigral iron levels compared to idiopathic PD patients and healthy controls (33), suggesting that iron dyshomeostasis could be an early and contributing feature in LRRK2-associated PD. However, the cellular and molecular mechanisms underlying these clinical observations remain unknown.

Autosomal dominant missense mutations in LRRK2 cause a late-onset form of familial PD while genome-wide association studies have identified risk-factor variants of LRRK2 linked to idiopathic PD (25). PD-linked mutations segregate within the kinase domain (e.g. G2019S) and the Roc-COR tandem domain (e.g. R1441C, Y1699C) of LRRK2 and increase phosphorylation of over a dozen Rab GTPase substrates at a conserved threonine/serine residue (34,35) disrupting intracellular trafficking and endolysosomal dynamics (26–29). Given that Rab GTPases govern critical steps in membrane trafficking and receptor recycling (36), these pathways offer a potential intersection between LRRK2 function and iron regulation. In our previous work, we observed alterations in trafficking of the transferrin receptor (TfR), that mediates iron uptake in most cell types (37–40), in mutant LRRK2 overexpression models (32). Importantly, we reported an increase in brain ferritin and iron load in homozygous G2019S KI mice compared to WT controls in acute pro-inflammatory conditions (32). Whether such disturbances occur basally and across different LRRK2 mutations, particularly in human neurons and glia when expressed in the heterozygous state, remains unclear. Astrocytes are key regulators of brain iron levels and availability and are implicated in PD neuropathology (41–47). Furthermore, LRRK2 expression is enriched in astrocytes and we and others have reported effects of LRRK2 mutations on astrocytic function (48,49). How endogenous LRRK2 mutations specifically affect cellular iron status and distribution in human astrocytes and neurons as well as the role of altered iron in neuronal damage remain unexplored.

In this study, we leverage patient-derived and isogenic induced pluripotent stem cell (iPSC) models to investigate how pathogenic LRRK2 mutations influence iron regulation in human neurons and astrocytes. By examining a wide panel of iron-related phenotypes, our data revealed convergent effects of heterozygous R1441C, Y1699C and G2019S LRRK2 mutations on intracellular and lysosomal iron content that were rescued by LRRK2 inhibition. Furthermore, we report distinct effects of LRRK2 mutations on ferritin levels and a concomitant effect on basal Iron Regulatory Protein Iron-Responsive-Element (IRE)-binding activity, suggesting alterations in ferritin expression and or post-transcriptional regulation. Pathogenic LRRK2 kinase activity is thought to inactivate Rab8a, which showed a special relationship with cellular iron. Genetic Rab8a deficiency recapitulated key iron-related defects observed in LRRK2 mutant cells, while exogenous Rab8a expression normalized lysosomal iron levels in heterozygous R1441C LRRK2 cells. Basal ROS levels and lipid peroxidation were elevated in LRRK2 mutant neurons and reduced by iron chelation or LRRK2 kinase inhibition. Importantly, kinase-dependent increases in lysosomal iron and lipid peroxidation were observed in LRRK2-mutant dopamine (DA) neurons, supporting the engagement of ferroptotic signaling in this cellular model of PD. Our data highlight a discrete pathway affected by pathogenic LRRK2 mutations that results in the misregulation of this transitional metal, leading to cellular injury. These data suggest a central role for iron dyshomeostasis in the interplay between LRRK2 signaling and ROS reported across many PD models and thought to contribute to PD (50–52), .

## Methods

### Induced pluripotent stem cell (iPSCs) cultures

We used iPSC lines carrying heterozygous LRRK2 mutations from three independent sources. BR33 were derived from a Caucasian donor, who was deeply phenotyped as part of the ROS/MAP longitudinal aging studies (53,54) and determined to not be cognitively impaired at death at age >89 and free from genetic variants that confer risk of PD. ND35371 (R1441C heterozygous) and ND50051 (G2019S heterozygous) were obtained by the NINDS Repository. Five G2019S heterozygous, two R1441G heterozygous and five WT iPSC lines were obtained from the Parkinson’s Progression Markers Initiative dataset (PPMI, ppmi-info.org) along with longitudinal clinical data. The isogenic mutant LRRK2 lines (all heterozygous) were a kind gift from Dr. Mark Cookson (NIA, NIH) (55).

### CRISPR/Cas9 Genome Editing of iPSCs and HEK293 cells

Rab8a and Rab10 knockout lines were generated in iPSCs and HEK293 cells as described before (56). Briefly, single guide RNAs (sgRNAs) were selected using a web-based design tool (http://crispr.mit.edu). Rab8a sgRNA: 5’ GAACTGGATTCGCAACATTG 3’; Rab10 sgRNA: 5’ ATGGCTTAGAAACATAGATG 3’. These were cloned into pXPR_003 (Addgene #52963) and sequenced using the primer 5′-GATACAAGGCTGTTAGAGAGATAATT-3′ to determine clones that successfully integrated the sgRNA. BR33 iPSCs were generated and characterized in collaboration with the New York Stem Cell Foundation (NYSCF) using described methods (54,57,58). iPSCs were co-transfected with plasmids that express the sgRNA plasmid and a nuclease-active SpCas9 expression vector (Addgene #62988). After 2 days, cells that were successfully transfected with the two plasmids were selected by puromycin treatment for 4 days. Individual clones were identified by plating ∼1 cell per well in a 96-well dish and allowed to grow for 2 weeks. Monoclonal lines were identified, expanded, sequenced, and stocked. Amplification of Rab8a and Rab10 genes was conducted according to manufacturer’s protocol (Invitrogen K2030-01) and homozygous gene editing was confirmed by Sanger sequencing. Multiple sequence alignment was performed using ClustalW in BioEdit.

### iPSC differentiation into mature astrocytes

iPSCs were differentiated into iAs using the protocol developed by Canals and colleagues (59). Briefly, iPSCs were maintained in mTeSR media and transduced with lentivirus to stably express pTet-O-NFIB (hygromycin), pTet-O-SOX9 (puromycin) and FUdelta GW-rtTA. Differentiation of pure human iA cultures was achieved by culturing transduced cells in doxycycline to induce expression of Sox9 and Nfib and to select against non-differentiated cells with selection media (puromycin and hygromycin). Human iAs were studied at day ∼21 post-differentiation. iAs show marked expression of GFAP by staining and immunoblot and robust expression of LRRK2 and relevant Rab GTPases.

### Differentiation of NGN2-induced neurons

iPSCs were differentiated into NGN2-induced neurons (iNs), as before (29,56,60). Briefly, iPSCs were cultured on Matrigel-coated plates in StemFlex (A33493) media and co-transduced with lentivirus packaged with pTet-O-NGN2-puro and Fudelta GW-rtTA plasmids (Zhang et al., 2013) for 2 days and passaged for expansion. NGN2-transduced iPSCs were thawed in StemFlex media with ROCK inhibitor (10 μM; Stemcell Technologies, 72304) for 24 hr then plated at 2 × 10^6^ cells/10 cm plate in StemFlex and grown until 75% confluent. For differentiation, on day 1 cells were fed with KnockOut media (Gibco 10829.018) supplemented with KnockOut Serum Replacement (Invitrogen 10928-028), 1% MEM non-essential amino acids (Invitrogen 11140), 1% GlutaMAX (Gibco 35050061) and 0.1% βME (Invitrogen 21985-023) (KSR) with doxycycline (2 μg/ml, Sigma, D9891-5g) to induce NGN2 expression. On day 2, they were fed with a 1:1 ratio of KSR:N2B media (DMEM F12 supplemented with 1% GlutaMAX, 3% dextrose and N2-Supplement B; StemCell Technologies 07156) with puromycin (5 μg/ml; Life Technologies, A11138-03) and doxycycline to select for transduced cells. On day 3, the cells were fed with N2B media with B27 (1:100; Life Technologies, 17504-044), puromycin, and doxycycline. On day 4, induced neurons (iNs) were frozen down in 10% DMSO/FBS in Neurobasal media (NBM Gibco 21103-049) supplemented with B27, BDNF (Peprotech, 450-02), CNTF (Peprotech, 450-13), and GDNF (Peprotech, 450-10) all at 100 ng/uL, ROCK inhibitor (10 μM), puromycin, and doxycycline. iNs were plated and grown in NBM with B27, BDNF, CNTF, GDNF, puromycin, and doxycycline until day 21.

### iPSC differentiation into DA neurons

iPSCs were cultured in StemFlex (A33493) media and co-transduced with lentivirus packaged with pTet-O-NGN2-puro and Fudelta GW-rtTA plasmids for 2 days and passaged for expansion, as before(29,56,60). To begin the dopaminergic neuron destination, NGN2-transduced iPSCs were thawed in mTeSR Plus media (StemCell technologies,100-0276) with 10 μM ROCK inhibitor (StemCell Technologies, 72304) and plated at 2 x 10^6^ cells/10 cm plate coated with Matrigel (Corning, 354277) and grown to 75% confluency - considered Day -1 (D-1). The iPSCs were differentiated into mature dopaminergic neurons using established protocols(61,62). Briefly, at D0 the cells were treated media containing growth factors and 2 μg/mL Doxycycline (ThermoFisher, J6057914) (Dox; present through differentiation) to begin the differentiation and Puromycin (ThermoFisher, A1113803) was added to the formulation for D1. On D2, cells were plated in media with containing growth factors and Dox with ROCK inhibitor at 5 x 10^4^ cells/well in 96 well plates (Griener, 655090) coated with Poly-L-ornithine hydrobromide (Sigma-Aldrich, P3655) - Laminin (Sigma-Aldrich, CC095-M). STEMdiff Midbrain Neuron Differentiation (Kit) media (StemCell Technologies, 100-0038) was used on D3 & D4 supplemented with Dox, 1 μM Cytarabine (Sigma-Aldrich, 251010), and 200 ng/mL SHH (StemCell Technologies, 78065). Daily media changes continue D5-D8 in the presence of Dox and SHH. D9-D21, cells were cultured using STEMdiff Midbrain Neuron Maturation (Kit) media (StemCell Technologies, 100-0038) with half media changes every third day.

### PPMI UPDRSIII data

Data used in the preparation of this article was obtained from the Parkinson’s Progression Markers Initiative (PPMI) database (www.ppmi-info.org/access-data-specimens/download-data), RRID:SCR_006431. Openly available UPDRSIII data for five G2019S and two R1441G mutations carriers were used (Tier 1 data downloaded between 03 November 2023 and 10 November 2023).

### IRE-binding activity assays

IRP activity was assessed using biotinylated IRE RNA probes, as previously described (63). Briefly, probes were 3’-biotinylated and sourced from Eurofins Genomics or IDT. Cell or tissue lysates were prepared in lysis buffer (40 mM KCl, 25 mM Tris-Cl pH 7.5, 1% Triton X-100) with protease inhibitors (ThermoFisher #78430), 1 mM DTT, and 20 U/ml RNase inhibitor (ThermoFisher #AM2696). For binding, 5 pmol of annealed IRE probes were incubated with 100 μg DynaBeads M280 for 20 min at room temperature, washed three times, then mixed with 100 μg lysates for another 20 min. IRP-bound complexes were isolated magnetically, washed, and resuspended in 100 μL LDS buffer, denatured at 95 °C for 5 min, and analyzed by western blot (25 μL on 4-12% Bis-Tris gels) for IRP1 and IRP2.

### ICP-MS

iPSCs in culture were collected in ice-cold PBS and divided equally into two tubes for ICP-MS and biochemical analysis of protein levels, pelleted and frozen in dry ice. Frozen iPSC pellets were resuspended in 200 μl PBS and mixed with 200 μl of concentrated trace-metal-grade nitric acid (Fisher) in 15 ml Falcon tube. The samples were digested at 85°C overnight, diluted to 4 ml with deionized water, and then analyzed by Agilent 7900 ICP-MS at the Analytical Toxicology Core Laboratory (ATCL) at the University of Florida. Metal levels were normalized to total protein content.

### Cell Treatments

For iron overload or chelation, cells were treated with 150 μM ferric ammonium citrate (FAC) (SIGMA) or 100 μM of deferoxamine in normal media overnight prior to analysis by imaging or western blotting. For rescue experiments, cells were treated with 100 nM MLi-2 (Tocris) for 2 hrs or 7 days in culture prior to analysis.

### iPSC transfection

Human iPSCs were transiently transfected with GeneJuice (Millipore) at ∼60-70% confluency using a 3:1 μL GeneJuice:μg DNA ratio prepared in Opti-MEM (ThermoFisher). DNA-GeneJuice complexes were added directly to cells in mTeSR1 medium, and transgene expression and cellular analyses were performed 24-48 hours post-transfection. The constructs eGFP-Rab8a (Addgene #86075) (64) and pcDNA3.1 eGFP (Addgene #129020) were used.

### Western Blot

Cells were lysed in cell lysis buffer (50 mM Tris-HCl, 150 mM NaCl, 0.5 mM EDTA, 0.5% (v/v) sodium deoxycholate, 1% (v/v) NP-40, pH 8) with protease and phosphatase inhibitors for 30 min. Lysates were centrifuged at 14,000 x *g* for 15 min at 4°C, and supernatants were quantified by BCA assay. Samples were prepared in 1x SDS-PAGE loading buffer, denatured at 65°C for 10 min, resolved on an SDS-polyacrylamide gel, and transferred to PVDF membrane. Membranes were blocked with 5% BSA (Sigma) prior to probing with primary antibodies against FTH1 (Cell Signaling 4393S), FTL (abcam AB69090), TfR1 (13-6800), DMT1 (abcam AB123085), Cyclophilin B (abcam AB16045). Secondary LiCOR antibodies were used for detection using the LiCOR model M scanner. Uncropped blots are presented in Supplementary Figure 10.

### Immunocytochemistry

Cells were seeded at 120,000/well (24-well plate) on 12 mm coverslips precoated with Geltrex basement membrane matrix (A1413201, Thermo) and cultured as described above. Cells were fixed in 4% (w/v) formaldehyde/PBS for 15 minutes, permeabilized in 0.2% Triton X-100/PBS for 10 minutes at RT, blocked in 5% (v/v) FBS in PBS, and incubated with primary antibodies (pT73 RAB10: MJF-R21-22–5, ab241060, Abcam; TH: CPC-TA, EnCor Bio) in 1% (v/v) FBS/PBS overnight at 4C. Following 3 washes in PBS, the cells were incubated for 1 hour with secondary antibodies (Alexa Fluor 488, 568, 647-conjugated; ThermoFisher). After 3 PBS washes, the coverslips were mounted, and the cells were imaged by confocal microscopy (Nikon CSU-W1-SORA).

### High-content imaging and quantitative analysis

For high-content live-cell imaging, cells were plated at 10,000 cells per well in 96-well black-wall clear-bottom plates (μClear, Greiner), labeled with FerroOrange (Dojindo), HMRhoNox-M (Lumiprobe), Liperfluo (Thermo), LysoTracker Red (Invitrogen) and CellROX (Thermo) according to the manufacturer’s specifications, and 20 ng/ml of Hoechst. CellLight Lamp1-emGFP BacMam 2.0 or LysoTracker (Thermo) were used to visualize lysosomes according to the manufacturer’s specifications. Labeled live cells were imaged using the Lionheart high-content automated microscope (Agilent) (10x or 20x objectives), or the ImageXpress HCS.ai high-content confocal imaging platform (Molecular Devices; 60x objective), collecting multiple fields per well for coverage of >800 cells per well. The DAPI, GFP, RFP/TRITC and Cy5 channels were used and images were analyzed on the Gen5 software (Lionheart; Agilent) or InCarta software (Molecular Devices). For experiments involving GFP control vector or GFP-Rab8a expression, quantitative analyses were restricted to GFP-positive cells to ensure consideration of successfully-transfected cells. To account for plate-to-plate variability in staining intensity, fluorescence measurements were normalized to the corresponding control condition within each imaging plate before pooling data across biological replicates.

### Quantification of lysosomal iron using whole-cell, lysosome-restricted segmentation

Quantitative analysis of HMRhoNox-M fluorescence was performed using the InCarta image analysis software (Molecular Devices) using two complementary segmentation strategies. For whole-cell analysis, nuclei were identified using Hoechst staining and used as primary objects. Cytoplasmic regions were generated based on the primary objects (nuclei) using TRITC staining for cell perimeter, producing a whole-cell mask for quantification of integrated HMRhoNox-M signal on a per-cell basis. For lysosome-restricted analysis, lysosomes were identified using LysoTracker staining. LysoTracker-positive puncta were segmented as subcellular objects using intensity-based thresholding and size filtering, producing a lysosomal mask per cell (similar to(65)), and HMRhoNox-M signal was quantified within lysosomes. Measurements were aggregated across multiple fields per well, with >800 cells analyzed per biological replicate.

### Imaris analysis

Following confocal microscopy, the Imaris platform (Oxford Instruments) was used to analyze the localization and distribution of lysosomal iron in lysosomes. Z-stack confocal images of iAs were processed through the Imaris Surface Contour module to render lysosomal iron, lysosomes and nuclear staining to surfaces and measure the distance between lysosomes and proximity to the nuclear edge throughout z planes in the 3D volume.

### Statistical Analyses

All experiments were conducted at least three independent times for three differentiations. Error bars indicate mean +SD. Statistical analysis was performed using GraphPad Prism software, using a one-way ANOVA with Dunnett’s post-hoc or two-way ANOVA with Tukey’s post-hoc test.

## Results

### LRRK2 mutations dysregulate iron-related pathways in patient-derived and isogenic iPSCs and neurons in a kinase-dependent manner

In our previous work, we identified a signature of altered iron pathways in the kidneys of homozygous G2019S knockin mice by unbiased proteomics, highlighting divergent expression of endolysosomal and mitochondrial factors involved in iron regulation *in vivo* (35). Separately, we reported increased accumulation of iron in the brains of homozygous G2019S LRRK2 mice following inflammation compared to WT controls (32). These data were the first to suggest that G2019S LRRK2 dysregulates aspects of intracellular iron homeostasis in LRRK2 models. To examine whether these pathways are affected by endogenous LRRK2 mutations beyond G2019S, and, when present in the heterozygous state most commonly associated with PD, we assessed different aspects of iron homeostasis in iPSCs derived from LRRK2-PD patients. We quantified cytosolic iron levels in six WT, six G2019S, one R1441C, and two R1441G LRRK2 iPSC lines (all heterozygous) by high-content imaging of the FerroOrange probe, a fluorescent indicator of labile iron. We observed an increase in the steady-state labile ferrous iron pool across different LRRK2 mutant lines compared to WT lines (Figure 1A, B). Each iPSC line is depicted in a different color to reflect the biological variability observed across the many patient-derived lines (Figure 1B). Increased total iron loading in the R1441C LRRK2 mutant iPSCs was further confirmed by ICP-MS (Figure 1C). Lastly, the small molecule LRRK2 kinase inhibitor MLi-2 reversed the increase in labile iron levels in heterozygous G2019S iPSC lines compared to WT, with a partial rescue in R1441C LRRK2 iPSCs (Figure 1D, E). Next, we investigated proteins involved in iron import (TfR1, DMT1) and iron storage (ferritin light chain; FTL). We observed an overall increase in FTL levels across the R1441C/G and G2019S mutant lines compared to WT iPSC controls (Figure 1F, G), while the levels of TfR1 and DMT1 were comparable between lines (Figure 1F and Supplementary Figure 1A, B). Despite an overall effect of mutations on iron-related readouts, we noted biological variability in the levels of labile iron (Figure 1B) and ferritin (Figure 1G) across the different patient-derived iPSC lines. We observed moderate-to-strong correlation between labile iron levels (FerroOrange) and FTL levels (R^2^=0.5660, F(1,13), p=0.0012) (Figure 1H), and a strong correlation between labile iron (FerroOrange) and total iron levels detected by ICP-MS (R2=0.8805. F(1,1), p=0.2247) (Figure 1I). We did not observe a correlation between ferritin levels in the iPSC lines and motor symptom severity (UPDRS III scores) in LRRK2 PD or control groups reported for the iPSC donors (Supplementary Figure 1C).

**Figure 1.**
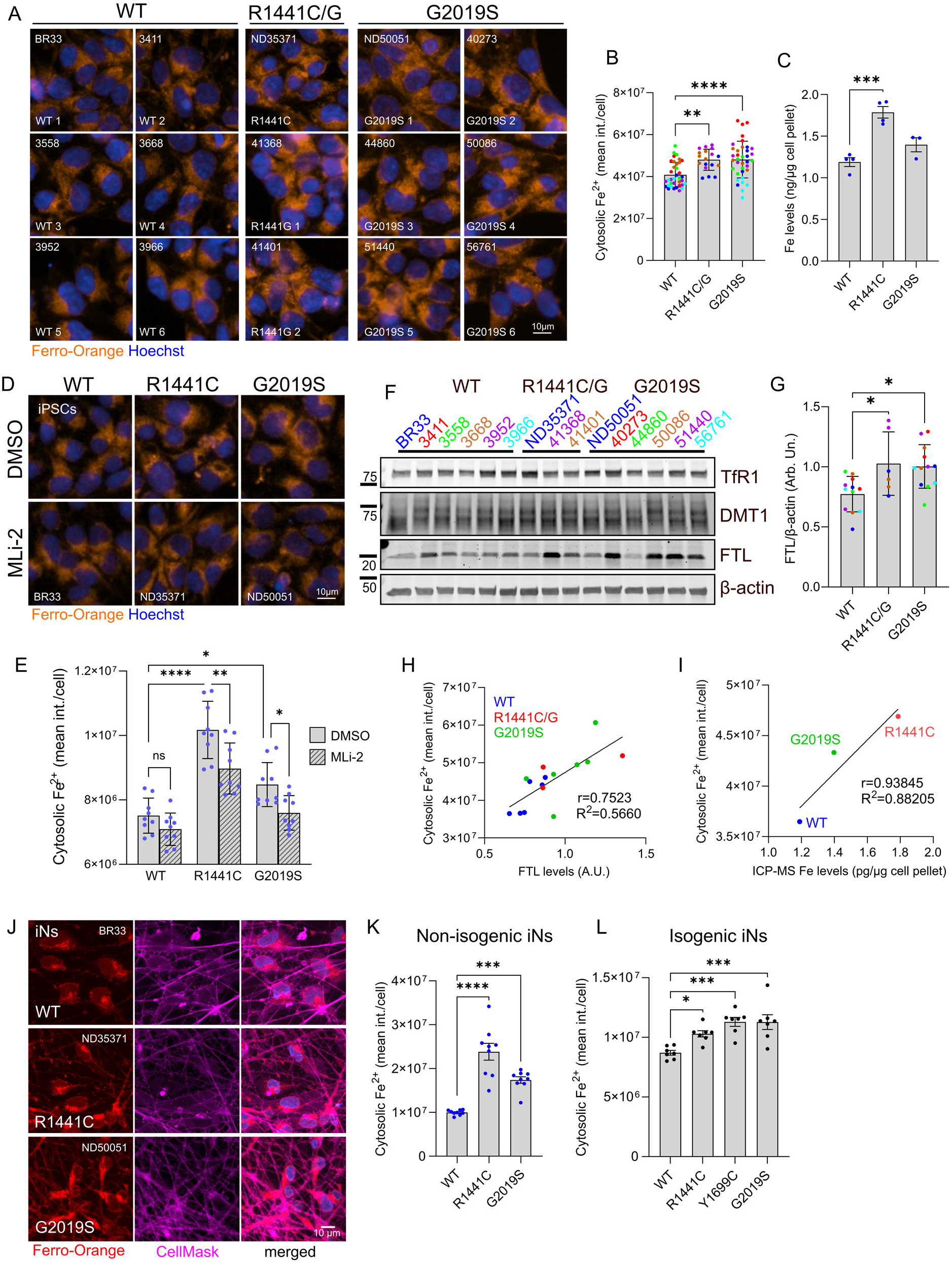
LRRK2 mutations dysregulate iron-related pathways in human iPSCs and iNs in a kinase-dependent manner. (A, B) Cytosolic free iron imaging and quantitation (FerroOrange) in six WT, six G2019S, one R1441C, and two R1441G LRRK2 iPSC lines (PPMI, NINDS Repository) (N=6 wells each line; each line represented in a different color within each genotype group; one-way ANOVA, Dunnett’s post-hoc pair-wise comparison). (C) Quantitation of total iron in WT, R1441C and G2019S iPSCs (NINDS Repository) by ICP-MS. (D, E) Imaging and quantitation of cytosolic free iron (FerroOrange) in mutant LRRK2 iPSCs (NINDS Repository) following Mli-2 treatment (100nM, 7 days) (two-way ANOVA, Tukey’s post-hoc; N=9 biological replicates (>800 cells per N), Genotype ****p=0.0001, Treatment ****p<0.0001, Interaction p=0.2318). (F, G) Western blot and quantitation of levels of iron-related factors in WT, heterozygous R1441C/G, and G2019S iPSCs (N=2 biological replicates per line; 6 WT, 2 R1441G, 1 R1441C and 6 G2019S lines, one-way ANOVA, Dunnett’s post-hoc; *p<0.05). (H) Scatter plot showing a positive correlation between cytosolic iron (FerroOrange) and FTL levels (r=0.7523). (I) Scatter plot showing a positive correlation between cytosolic ferrous iron levels (FerroOrange) and total cellular iron quantified by ICP-MS across iPSC lines. Each data point represents the mean value for a single iPSC line (WT: BR33, R1441C: ND35371, G2019S: ND50051), calculated from >4 independent biological replicates for ICP-MS and 6 independent imaging wells for FerroOrange (>800 cells per well). Pearson correlation shown. (r=0.93845). (J to L) Cytosolic iron imaging and quantitation in mature iNs carrying heterozygous LRRK2 mutations (J, K) as well as isogenic iNs versus controls (NIA, NIH) (L) (one-way ANOVA, Dunnett’s post-hoc, N=8 biological replicates, (>800 cells per N), *p<0.05, **p<0.01, ***p<0.001, ****p<0.0001).

Since iron dyshomeostasis is a key feature in PD pathology and our data support altered iron levels in LRRK2 mutant iPSCs, we next examined human iPSC-derived NGN2-induced neurons (iNs) to understand how this change translates in post-mitotic cells. iPSCs (NINDS Repository) were differentiated into iNs by forced expression of NGN2 via lentiviral gene delivery, according to established protocols (29,56,60) (Supplementary Figure 2). We detected an increase in the labile iron pool across heterozygous R1441C and G2019S LRRK2 neurons compared to WT controls (Figure 1J, K). Considering the biological variability observed in these patient-derived iPSC lines (Figure 1B), we next tested this phenotype in a series of isogenic LRRK2 mutant iPSC lines. These lines have been generated by CRISPR/Cas9 in a single female iPSC background, A18945, with editing validated and clones subsequently characterized for morphology, karyotype and differentiation potential(55). NGN2-induced neurons generated from this isogenic LRRK2 series validated the phenotype with an increase in the labile iron pool across R1441C, Y1699C and G2019S heterozygous mutant iNs, compared to WT controls (Figure 1L), supporting a generalizable effect of pathogenic LRRK2 mutations on iron homeostasis.

### LRRK2 mutations cause kinase-dependent accumulation of lysosomal iron in neurons and astrocytes

The FerroOrange iron probe is reported to detect free ferrous iron in the cytosol as well as in different organelles including ER, mitochondria and endolysosomes (66–68) (Supplementary Figure 3A). Thus, the data in Figure 1 do not differentiate between cytosol and specific subcellular compartments. Given the role of lysosomes as important sites for iron mobilization regulating intracellular iron availability (69–71), and the well-established role of LRRK2 in lysosome-related pathways reported by us and others (28,29,32,72–74), we focused on evaluating lysosomal iron in LRRK2 mutant cells.

We thus employed a reported lysosomal specific ferrous iron (Fe^2+^) probe (HMRhoNox-M) (75) to assess relative lysosomal Fe^2+^ levels across iPSCs, NGN2-induced neurons (iNs) and induced-astrocytes (iAs) (59) using high-content imaging and super-resolution microscopy (Figure 2). iAs were generated by viral delivery of doxycycline-inducible expression of SOX9 and NFIB (59). The expression of astrocytic markers and LRRK2 was confirmed at ∼21 days post-induction *in vitro* (Supplementary Figure 2). We first validated the lysosomal specificity of HMRhoNox-M by comparing its staining pattern to Lamp1-emGFP localization (Figure 2A). In both iPSC-derived human neurons and astrocytes (Figure 2A), HMRhoNox-M signal was enriched within Lamp1-emGFP positive structures, supporting its preferential localization to lysosomes. Quantification of HMRhoNox-M signal in iPSCs carrying heterozygous LRRK2 mutations revealed an increase in R1441C, Y1699C and G2019S LRRK2 mutant lines compared to isogenic WT controls (Figure 2B, C). The LRRK2 mutation-driven increase in lysosomal iron as detected by HMRhoNox-M was likewise observed in differentiated iNs (Figure 2D, E) and astrocytes (Figure 2F, G) generated from these parental isogenic iPSCs.

**Figure 2.**
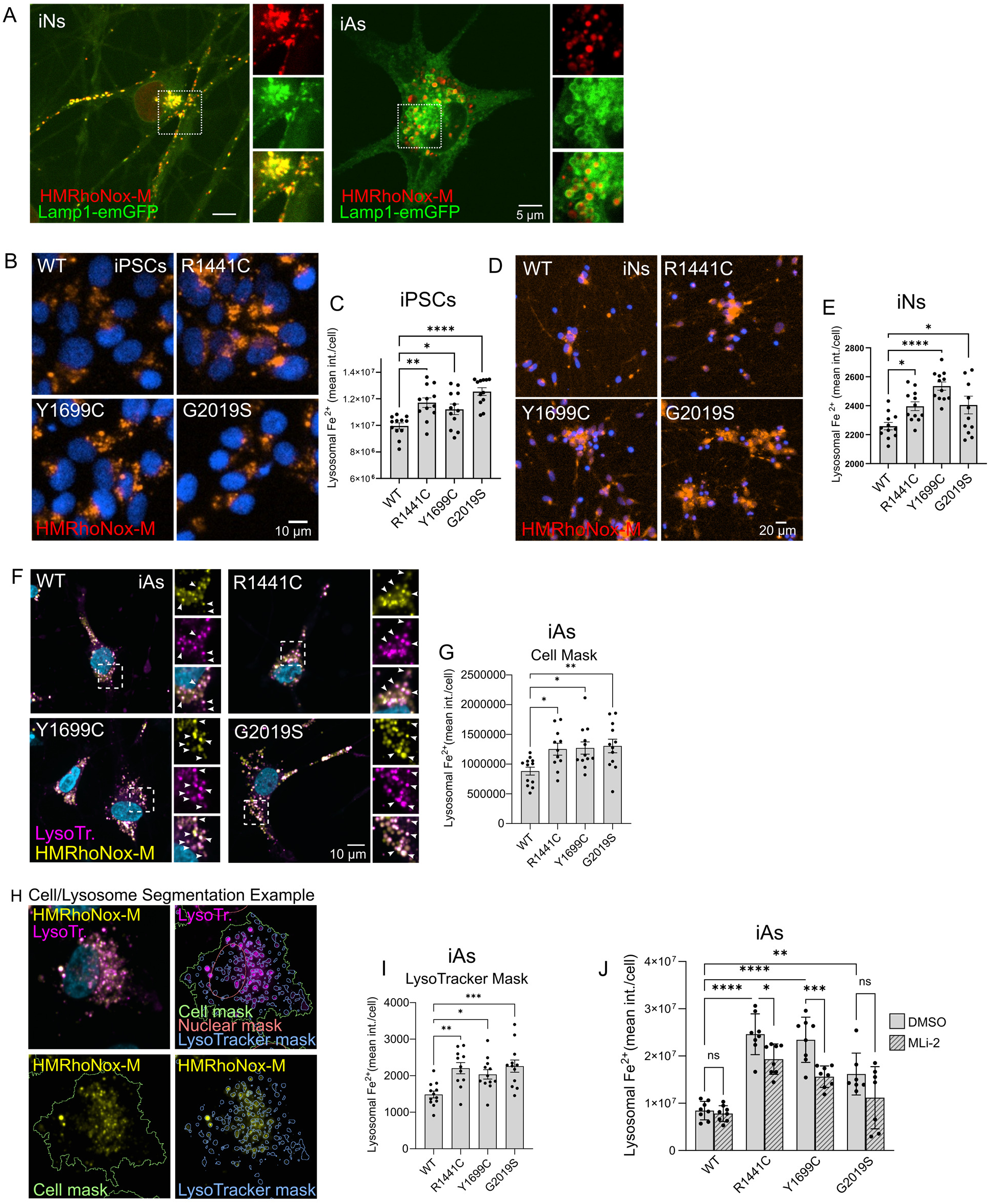
LRRK2 mutations increase lysosomal iron in isogenic neurons and astrocytes. (A). Representative images of lysosomal iron (HMRhoNox-M) and Lamp1-emGFP by super-resolution confocal microscopy in iNs and iAs. Lysosomal iron imaging and quantitation in heterozygous LRRK2 mutant isogenic iPSCs (B, C), isogenic iNs (D, E), and isogenic iAs (F, G). Quantification was performed using a whole-cell mask to capture total cellular HMRhoNox-M signal (one-way ANOVA, Dunnett’s post-hoc, N=8 biological replicates, (>800 cells per N), *p<0.05, **p<0.01, ***p<0.001, ****p<0.0001). (H) Representative images illustrating LysoTracker-based segmentation of acidic lysosomal compartments and corresponding detection of lysosomal iron using HMRhoNox-M. Cell, nuclear, and LysoTracker masks used for analysis are shown. (I) Quantification of HMRhoNox-M signal specifically within LysoTracker-segmented lysosomes in isogenic iAs, demonstrating genotype-dependent increases in lysosomal iron that recapitulate those observed using whole-cell mask-based quantification. Data are shown as mean ± SEM (one-way ANOVA with Dunnett’s post hoc test). (J) Lysosomal iron quantitation in iAs following treatment with MLi-2 in culture (100nM, 7 days) (two-way ANOVA, Tukey’s post-hoc; N=8 biological replicates (>800 cells per N), Genotype ****p=0.0001, Treatment ****p<0.0001, Interaction p=0.0844).

Notably, not all HMRhoNox-M-positive puncta overlapped with Lamp1-emGFP-labeled structures, particularly in astrocytes (Figure 2A), which could reflect incomplete labeling of acidic lysosomal compartments by the Lamp1 fusion protein (76). To address this possibility directly, we performed complementary analyses using LysoTracker, which labels acidic lysosomes independently of exogenous Lamp1 expression (Figure 2F; Supplementary Figure 3). Using this approach, HMRhoNox-M showed strong colocalization with LysoTracker-positive compartments (Supplementary Figure 3A). Quantification of HMRhoNox-M selectively within LysoTracker-segmented lysosomes (Figure 2H, I) recapitulated the genotype-dependent increases observed using a whole-cell mask-based analysis (Figure 2G, H). Finally, to exclude the possibility that differences in lysosomal Fe^2+^ levels reflected changes in lysosome abundance or morphology rather than iron content per organelle, we quantified lysosome number and size across the isogenic lines. No significant differences in these parameters were observed in heterozygous LRRK2 mutant iPSCs (Supplementary Figure 3B-E), indicating that elevated HMRhoNox-M signal reflects increased lysosomal iron load rather than altered lysosomal biogenesis or expansion.

Given that LRRK2 can localize to lysosomal membranes and that lysosomal position has been reported to correlate with LRRK2 kinase activity toward Rab GTPases (77), we next examined whether iron accumulation exhibited spatial heterogeneity within the lysosomal network. Quantitative analysis revealed that the increase in lysosomal iron content in R1441C iAs was more pronounced in the perinuclear compared to the peripheral lysosomes and that this increased iron load was preferentially associated with smaller lysosomes (< 3 µm^3^) (Supplementary Figure 3F, G, H). These data are consistent with perinuclear lysosomes being sites of active LRRK2 signaling (77), as demonstrated by work from us and others showing LRRK2-dependent effects on lysosomal function and proteolytic activity (26,56,78,79).

To determine whether the increase in lysosomal iron observed in LRRK2 mutant iAs is dependent on LRRK2 kinase activity, iAs were treated with MLi-2 for 7 days prior to analysis (Figure 2J). LRRK2 inhibition reduced lysosomal iron levels in all mutant lines but not WT, with partial rescue in the R1441C and Y1699C lines and complete rescue in the G2019S line (Figure 2J). Effective inhibition of endogenous LRRK2 kinase activity was validated in parallel iPSC cultures by assessing Rab10 phosphorylation at Threonine 73, which was elevated in R1441C iPSCs and robustly reduced following acute MLi-2 (2 hrs) treatment (Supplementary Figure 4A, B). To assess the selectivity of MLi-2 on LRRK2-mediated effects on iron homeostasis, LRRK2 KO iPSCs were treated with vehicle or MLi-2 for either acute (2 hrs) or prolonged (7 day) durations and cytosolic and lysosomal iron levels were assessed. MLi-2 had no effect on iron levels in LRRK2 KO cells (Supplementary Figure 4C-F), indicating that MLi-2 as employed here does not possess off-target effects with respect to iron homeostasis. Together, these results provide evidence that lysosomal iron levels are increased by a series of pathogenic LRRK2 mutations in a kinase-dependent manner.

### Ferritin levels are dysregulated in isogenic LRRK2 mutant iPSCs

Altered cytosolic and lysosomal iron demonstrate an impairment in how cells regulate their iron levels and distribution, so we further evaluated expression of key proteins involved in these processes. Iron uptake, transport, and storage are finely regulated at the post-transcriptional level by the iron regulatory protein (IRP) and iron-responsive element (IRE) signaling pathways (80). IRP1 and IRP2 are RNA-binding proteins that interact with IREs located within the untranslated regions of specific mRNAs, to modulate iron homeostasis. These interactions repress ferritin (both FTH1 and FTL) and FPN mRNA translation while stabilizing TfR mRNA (81–83). In proliferating iPSCs cultured without iron supplements, we observed an increase in total FTH1 and FTL levels in G2019S LRRK2 cells compared to isogenic WT controls, while R1441C LRRK2 iPSCs showed a trend of lower FTH1 (Figure 3A, B, C). Consistent with an IRP-mediated loss of translational repression of ferritin heavy and light chains, G2019S iPSCs also had lower basal TfR1 levels as compared to WT (Figure 3A, D). We therefore directly measured IRP1 and IRP2 levels and activity in these LRRK2 mutants. We observed a trend of downregulation of IRP1 in the ROC-COR domain mutants that reached significance in Y1699C iPSCs, supporting impairment in IRP1 signaling (Figure 3A, E). Separately, the G2019S iPSCs showed lower IRP2 expression levels with the ROC-COR domain mutants showing a trend toward decrease (Figure 3F). To evaluate IRE-binding activities of IRP1 and IRP2, we used a biotinylated-IRE probe to separate IRE-binding IRPs from cell lysates and analyzed by immunoblotting, as before (63). The IRE-binding activity of IRP1 was lower in G2019S iPSCs compared to the isogenic WT (Figure 3G), as were IRP2 protein levels and IRE-binding activity (Figure 3A).

**Figure 3.**
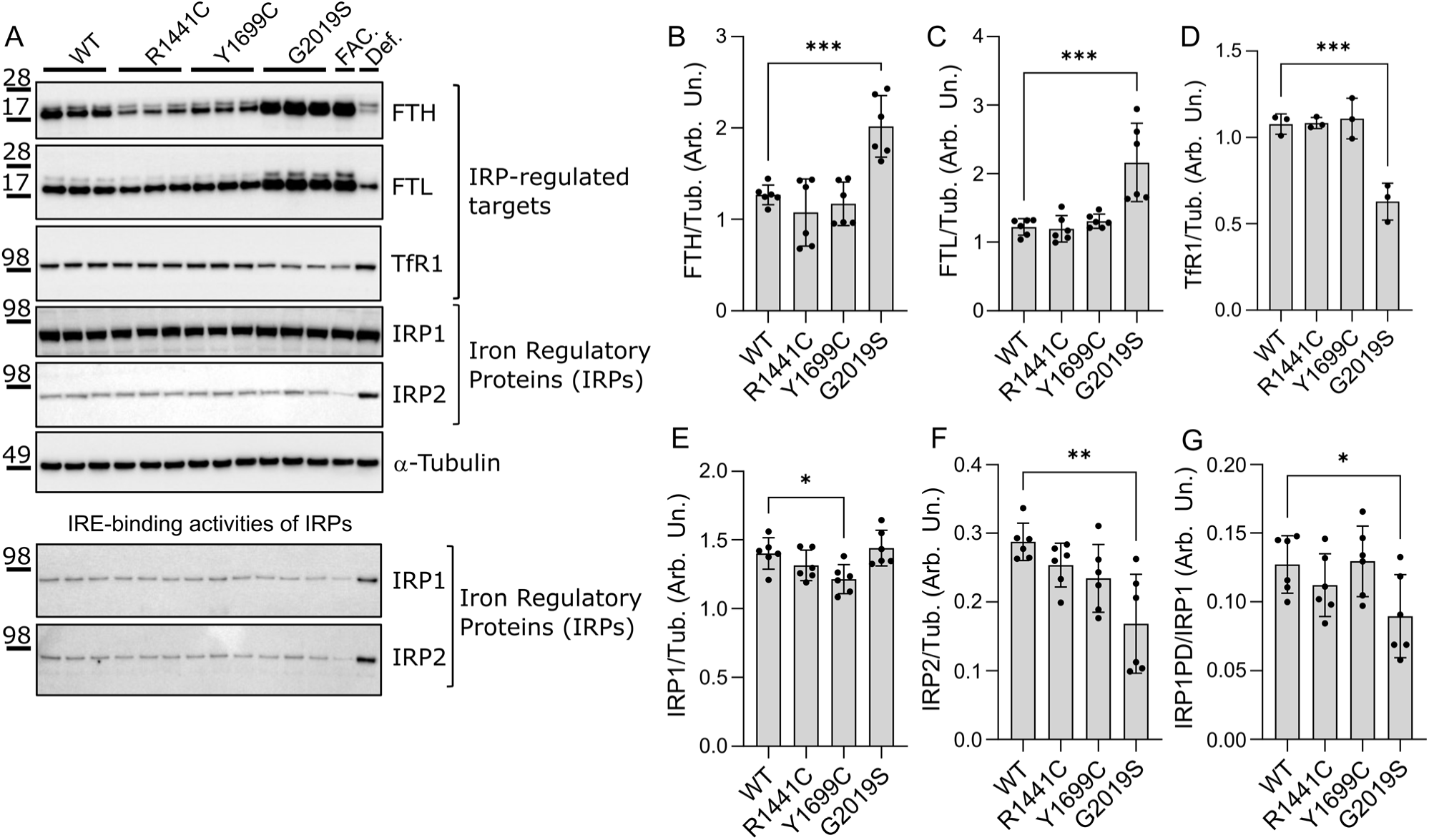
Ferritin heavy chain is dysregulated in isogenic LRRK2 mutant iPSCs. (A-F) Western blot analysis of IRPs (IRP1, IRP2) and IRP-regulated proteins (FTH1, FTL, TfR1) in isogenic LRRK2 mutant iPSCs. (A bottom panel, G) Western blot of IRE-bound IRP1 in isogenic LRRK2 mutant iPSCs and quantitation of IRE-binding. (one-way ANOVA, Dunnett’s post-hoc, N=6 biological replicates, *p<0.05, **p<0.01, ***p<0.001).

IRP1 activity is regulated by intracellular iron levels but can also be influenced by Fe-S cluster biogenesis as it requires a 4Fe-4S cluster to function as a cytosolic aconitase (81,84). Loss of the Fe-S cluster, under iron deficient conditions, shifts IRP1 to its IRE-binding form. In contrast, IRP2, does not coordinate an Fe-S cluster, and cellular iron availability is directly reflected by IRP2 protein levels, which increase when iron is scarce and decrease through degradation when iron is sufficient. To evaluate a potential effect of iron dyshomeostasis on mitochondrial function, we assayed levels of mitochondrial Fe-S-containing subunits (Supplementary Figure 5), noting no differences in basal levels of UQCRFS1 (Fe-S subunit of complex III), and SDHB (Fe-S subunit of complex II). While our data suggests a mild effect of LRRK2 mutations on IRP IRE-binding activities, we observe significant effects on ferritin levels, suggesting that separate mechanisms on ferritin homeostasis may be at play.

### LRRK2-Rab8a signaling controls cellular iron homeostasis

Phosphorylation of Rab GTPases by LRRK2 locks the molecules in an inactive state affecting their function (32,36,56,85). We previously showed that LRRK2-driven phosphorylation of Rab8a impairs its function in TfR trafficking, linking LRRK2 to iron homeostasis (32). Furthermore, we demonstrated that Rab8a and Rab10 knockouts (KO) produce distinct phenotypes in lysosomal integrity and PD-pathology related proteostasis (56). To test whether genetic inactivation of either of these Rab proteins is sufficient to drive dysregulation of cellular iron levels, we used CRISPR/Cas9 gene editing to generate Rab8a KO and Rab10 KO iPSCs (56) and HEK293 cells (Supplementary Figure 6), and assayed cellular iron load alongside matched isogenic controls. Using the FerroOrange probe to quantify labile cytosolic Fe^2+^ iron, we found that Rab8a but not Rab10 KO iNs showed an increase in free iron compared to isogenic WT controls, phenocopying the increase observed in mutant LRRK2 cells (Figure 4A. B). To determine whether Rab8a deficiency is causative for this phenotype, we next tested whether re-expression of Rab8a could restore iron homeostasis. Expression of exogenous GFP-Rab8a in Rab8a KO iPSCs significantly reduced FerroOrange signal, normalizing labile cytosolic Fe^2+^ iron levels toward those observed in WT cells (Figure 4C, D). These findings demonstrate that Rab8a loss is sufficient to increase labile cytosolic Fe^2+^ iron levels and that this phenotype is reversible upon Rab8a reintroduction. Notably, this increase in labile iron was specific to Rab8a loss, as Rab10 deficiency resulted in a divergent phenotype in iPSCs and HEK293 cells, supporting non-redundant roles for LRRK2-regulated Rab GTPases in iron homeostasis (Supplementary Figure 7).

**Figure 4.**
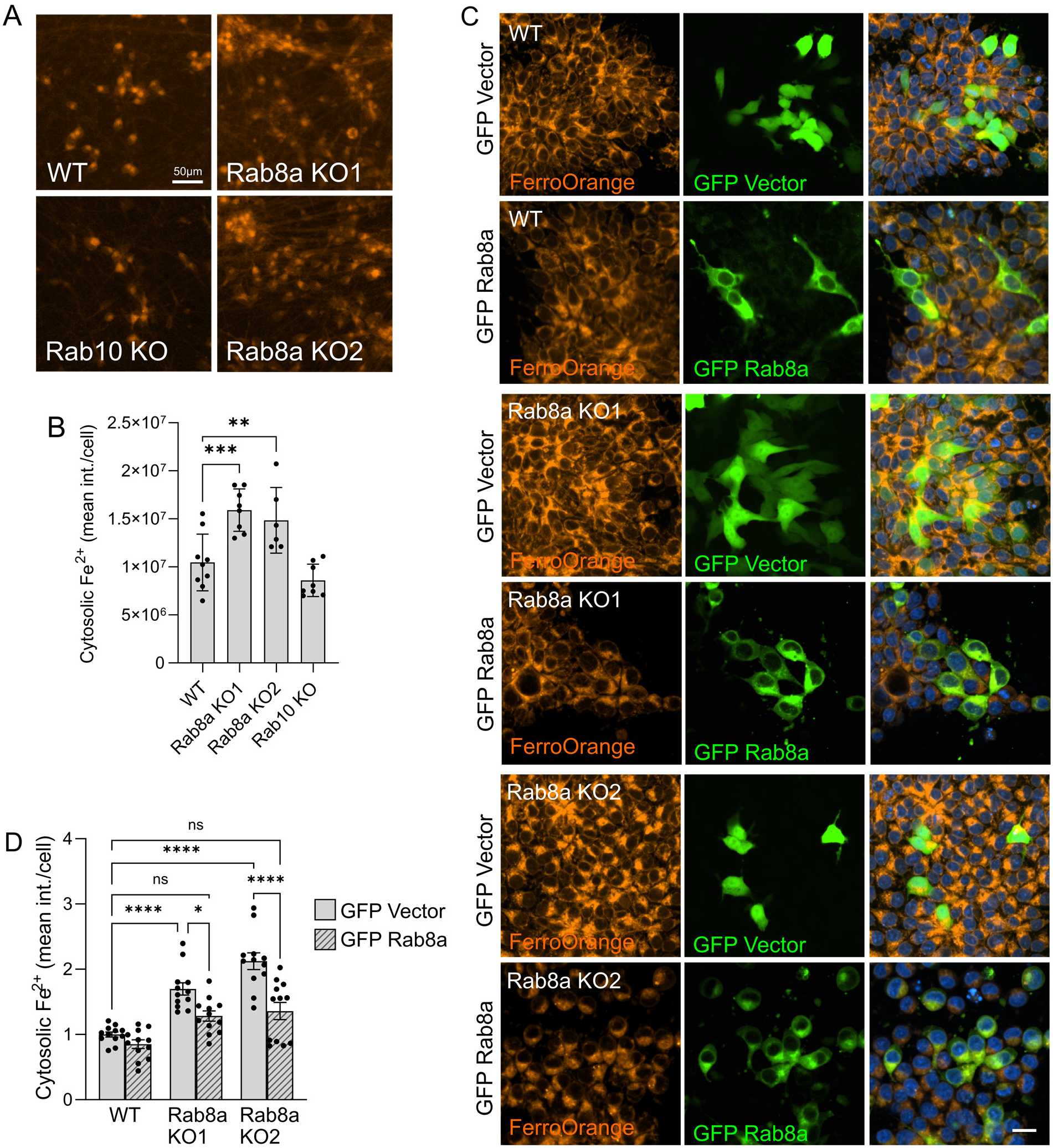
Knockout of the LRRK2 substrate Rab8a dysregulates cellular iron. (A, B) Imaging of cytosolic free Fe^2+^ iron and quantitation of iron levels in isogenic WT, Rab8a KO and Rab10 KO iNs by high-content imaging (one-way ANOVA, Dunnett’s post-hoc, N>6 biological replicates, *p<0.05, **p<0.01, ***p<0.001, ****p<0.0001). (C, D) Representative images and quantitation of free cytosolic iron staining in WT and Rab8a KO iPSCs expressing either GFP control vector or GFP-Rab8a (two-way ANOVA, Tukey post-hoc; N=12 biological replicates (>800 cells per N), Genotype ****p<0.0001, cDNA ****p<0.0001, Interaction **p=0.0073).

To further assess how Rab8a and Rab10 deficiency influence cellular iron handling, we examined iron-related protein expression in Rab8a and Rab10 KO HEK293 cells under basal conditions and following overnight exposure to 150 µM ferric ammonium citrate (FAC). Rab8a KO cells exhibited elevated basal FTH1 levels compared to WT and Rab10 KO cells but showed a blunted FTH1 induction following FAC treatment (Supplementary Figure 7). In contrast, Rab10 KO cells displayed a robust increase in FTH1 upon iron overload. Basal TfR1 levels were increased in Rab10 KO cells, while both Rab KO lines exhibited a greater reduction in TfR1 expression following FAC treatment relative to WT. Consistent with these protein changes, Rab8a KO cells accumulated higher levels of labile iron under iron overload conditions, which was reversed by iron chelation. Together, these data further support Rab8a loss as a key driver of cellular iron accumulation; while highlighting that not all LRRK2-regulated Rab GTPases exert equivalent effects on iron-handling pathways.

The LRRK2-dependent increase in lysosomal Fe^2+^ iron was the most consistent phenotype across all pathogenic mutations and cell types analyzed. Since LRRK2 mutations are hypothesized to inactivate Rab substrates, we examined whether exogenous Rab8a over-expression could overcome this effect in R1441C LRRK2 mutant iPSCs. WT and R1441C iPSCs were transiently transfected to express either GFP or GFP-Rab8a and lysosomal iron was quantified in positively transfected cells. GFP-Rab8a expression in WT iPSCs resulted in a minor reduction in iron that did not reach statistically significant deviation from the GFP control. R1441C iPSCs transfected with GFP showed increased lysosomal iron compared to WT cells, as expected. However, expression of GFP-Rab8a in R1441C iPSCs significantly reduced lysosomal iron levels to those comparable to WT (Figure 5A-B). Collectively, these data identify Rab8a as a critical downstream effector of LRRK2-dependent iron dyshomeostasis and demonstrate that restoration of Rab8a activity is sufficient to rescue iron phenotypes associated with LRRK2 mutations.

**Figure 5.**
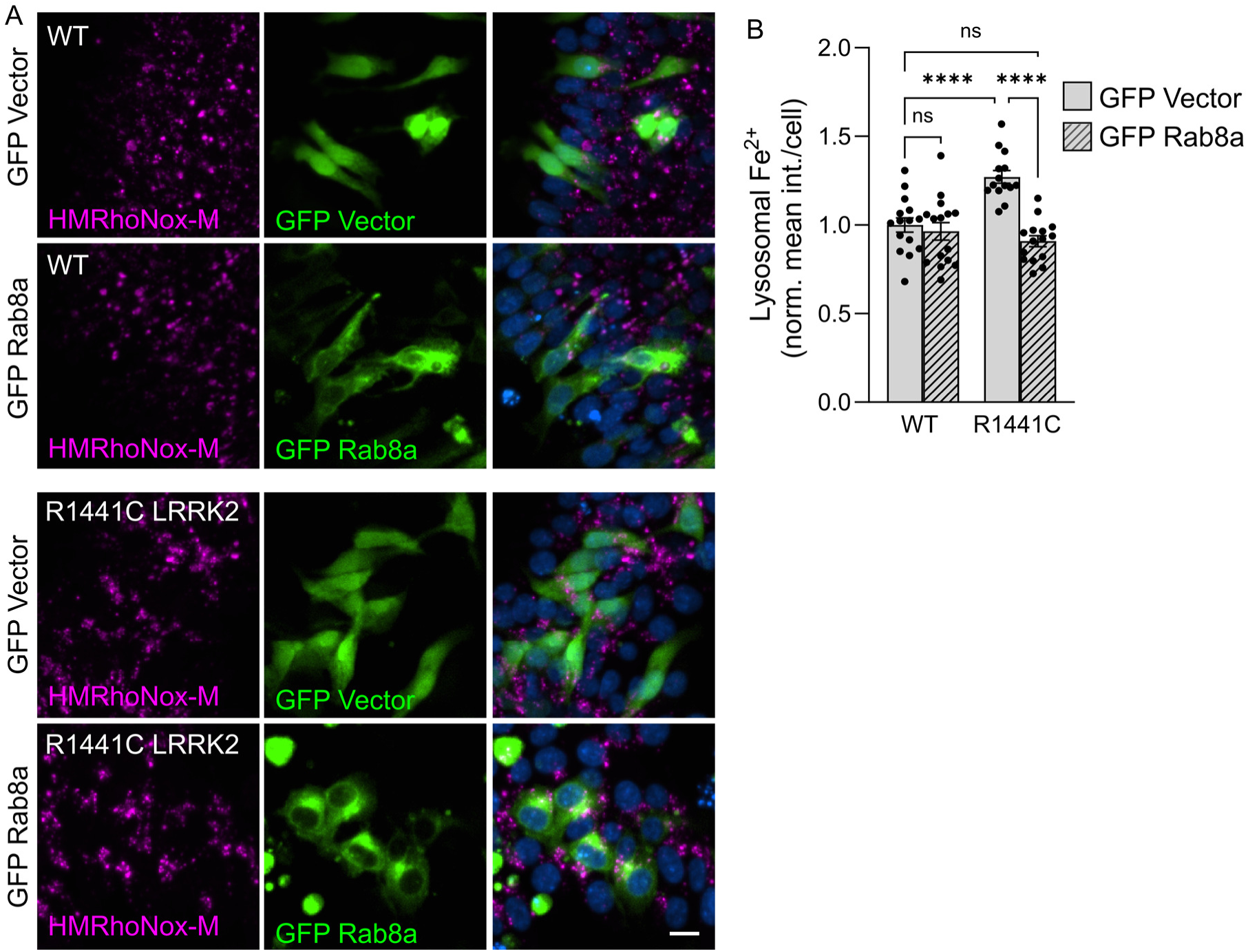
Exogenous Rab8a expression normalizes lysosomal iron levels in R1441C LRRK2 iPSCs. (A, B) Representative images and quantitation of lysosomal iron in WT and R1441C LRRK2 iPSCs expressing either GFP control vector or GFP-Rab8a. (two-way ANOVA, Tukey post-hoc; N=15 biological replicates (>800 cells per N), Genotype p=0.0096, cDNA ****p<0.0001, Interaction ***p=0.0002).

### Iron chelation and LRRK2 inhibition protect against LRRK2-driven oxidative stress and lipid peroxidation

Ferroptosis, the process of iron-mediated lipid peroxidation and cell death, is an emerging pathogenetic mechanism of interest in the neurodegeneration in PD (20,86–88). Iron catalyzes the conversion of hydrogen peroxide to more reactive free radicals via the Fenton reaction thereby promoting oxidative injury. Ferroptosis in neurodegeneration involves the simultaneous accumulation of brain iron and lipid peroxidation, which trigger a cascade of pathologic events including inflammation, myelin sheath degeneration, glial dysregulation and cell death (89). LRRK2 mutations have been linked to increased ROS (50,51) and ferroptosis signaling (90,91) in different model systems but the mechanism by which LRRK2 signaling leads to increased cell stress remains largely unclear. Here, we examined whether dysregulated lysosomal iron in LRRK2 mutant neurons contributes to oxidative stress and downstream lipid peroxidation. Compared to isogenic WT controls, all three LRRK2 mutations led to increased lysosomal iron in differentiated iNs (Figure 6A, B). As expected, the iron chelator deferoxamine (100 µM; 16 hours) significantly reduced lysosomal iron levels below control in all cells (Figure 6A, B). Assessment of cytoplasmic ROS using CellROX revealed a genotype-dependent increase in oxidative stress in heterozygous LRRK2 mutant neurons, which was reduced by deferoxamine treatment, consistent with its ability to chelate iron in the media (Figure 6C, D). We similarly assessed lipid peroxidation with Liperfluo, which produces a prominent punctate, vesicular pattern by confocal microscopy (Supplementary Figure 8A) (92), and used erastin, a ferroptosis inducer as a positive control (Supplementary Figure 8B, C) (93–96). All three pathogenic mutations in LRRK2 significantly increased lipid peroxidation levels in iNs, which was likewise attenuated by iron chelation with deferoxamine (Figure 6E, F). Similarly, at basal conditions in proliferating cells, we detected an increase in lipid peroxidation in R1441C and G2019S iPSCs compared to isogenic WT controls (Supplementary Figure 8B, C). These data are in line with recent work that places LRRK2 in a ferroptosis pathway and highlight lipid peroxidation as a relevant mechanism in LRRK2 signaling (90,91). Together, these findings demonstrate that pathogenic LRRK2 mutations promote iron-dependent oxidative stress and lipid peroxidation in neurons, consistent with a pro-ferroptosis phenotype.

**Figure 6.**
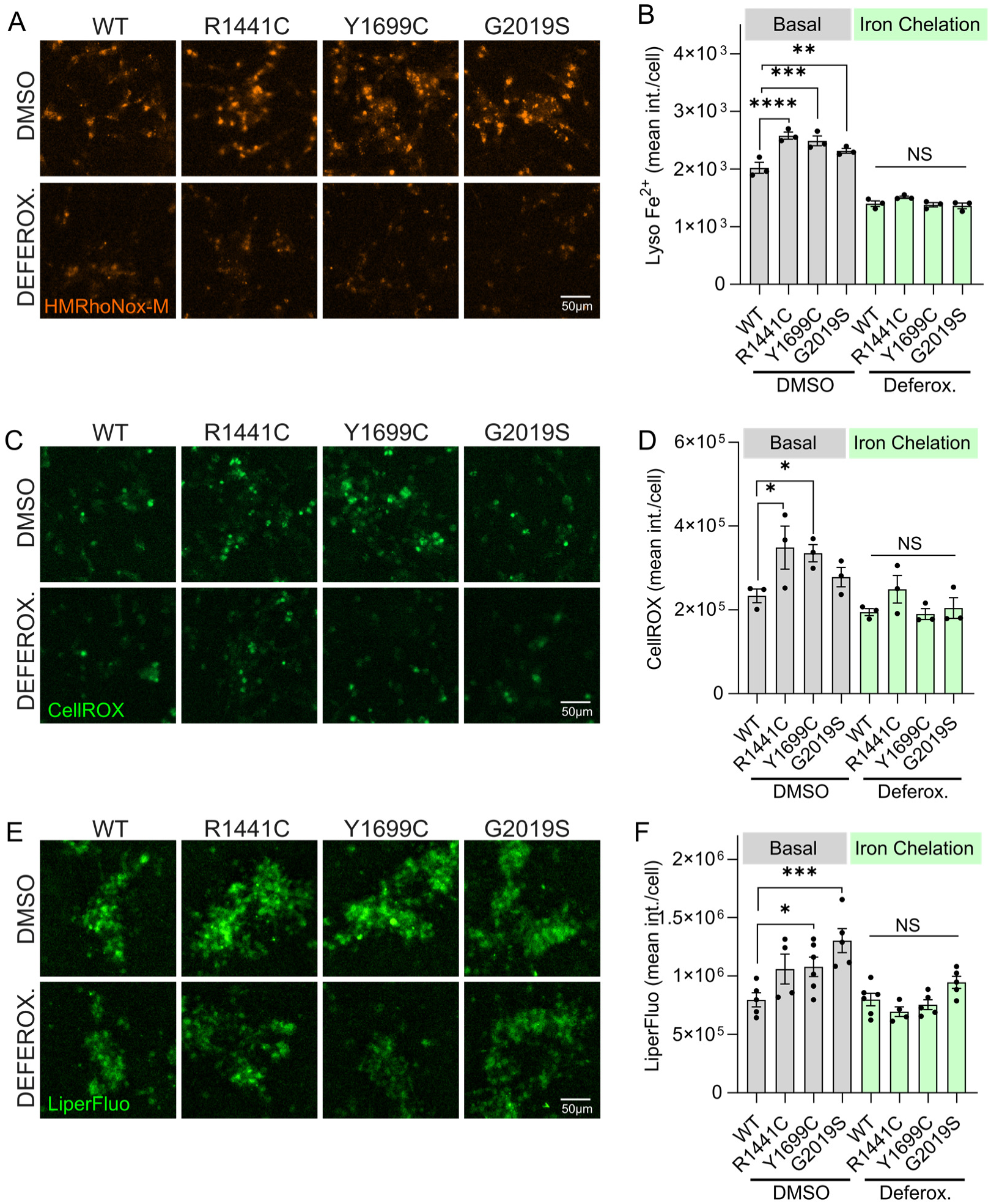
Iron chelation protects against LRRK2-driven ROS accumulation and ferroptosis. (A, B) High-content imaging and quantitation of lysosomal iron content (HMRhoNox-M) in iNs carrying heterozygous LRRK2 mutations versus isogenic WT controls, following treatment with 100 µM deferoxamine (16 hours) in normal media. (C, D) High-content imaging and quantitation of CellROX dye in LRRK2 mutant iNs. (E, F) Lipid peroxidation imaging (Liperfluo) and quantitation in iNs. (HMRhoNox-M: two-way ANOVA, Tukey post-hoc; N=3 biological replicates (>800 cells per N), Genotype ***p=0.0003, Treatment ****p<0.0001, Interaction **p=0.0034; CellROX: two-way ANOVA, Tukey post-hoc; N=3 biological replicates (>800 cells per N), Genotype *p=0.02, Treatment ***p=0.0002, Interaction p=0.3923; Liperfluo: two-way ANOVA, Tukey post-hoc; N=3 biological replicates (>800 cells per N), Genotype ***p=0.0009, Treatment ****p<0.0001, Interaction *p=0.0497).

To determine whether LRRK2 kinase activity contributes directly to lipid peroxidation, we next assessed the effect of pharmacological LRRK2 inhibition using MLi-2. To assess the selectivity of this inhibitor, LRRK2 KO iPSCs were treated with vehicle or MLi-2 (100 nM) for 7 days, with fresh inhibitor applied every 48 hr. No differences in basal Liperfluo signal were observed, confirming the selectivity of MLi-2 (Supplementary Figure 8D, E). We then examined the effect of selective LRRK2 kinase inhibition on lipid peroxidation in LRRK2 mutant iNs. Treatment of mature iNs with MLi-2 significantly reduced Liperfluo signal in R1441C, Y1699C, and G2019S neurons, reducing lipid peroxidation levels toward those observed in WT controls (Figure 7A, B). These data indicate that lipid peroxidation is increased in LRRK2 mutant neurons in a kinase-dependent manner.

**Figure 7.**
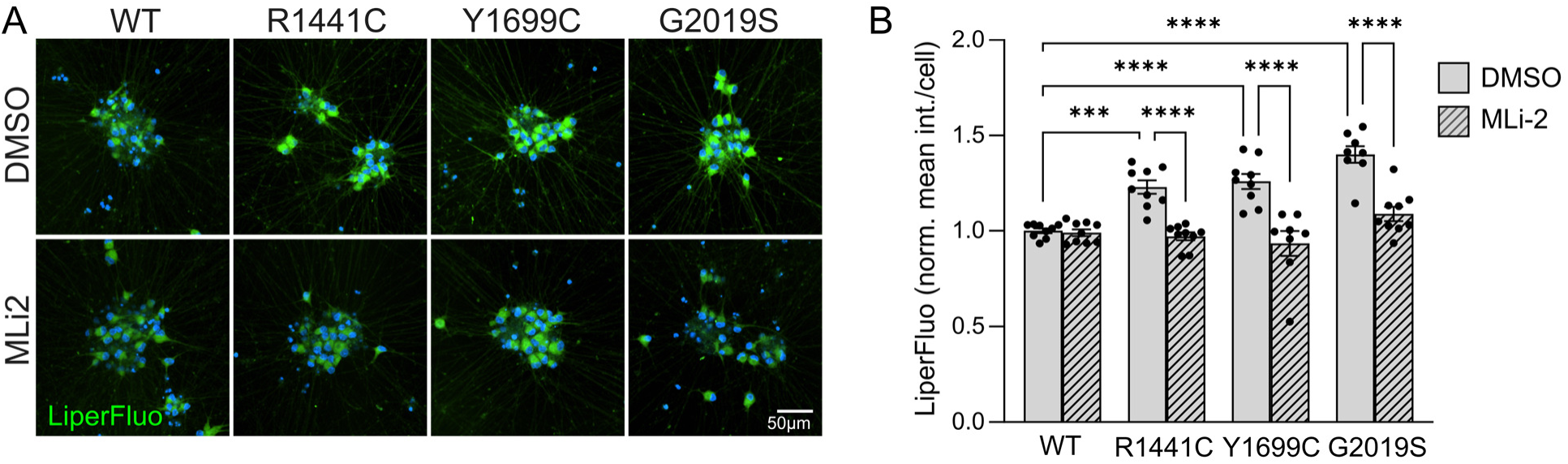
Selective kinase inhibition reduces lipid peroxidation in LRRK2-mutant neurons. (A, B) Representative images and quantitation of lipid peroxidation detected by Liperfluo in mature iNs expressing WT or isogenic LRRK2 variants following treatment with vehicle or MLi-2 (100 nM, 7 days) (two-way ANOVA, Tukey post-hoc; N=9 biological replicates (>800 cells per N), Genotype p<0.0001, Treatment p<0.0001, Interaction p<0.0001).

### LRRK2 kinase-dependent iron dysregulation in human DA neurons

Given the selective vulnerability of DA neurons in PD, we next examined whether LRRK2 mutations dysregulate iron homeostasis and oxidative stress in human iPSC-derived DAs. DA neurons were generated from isogenic WT and LRRK2 mutant iPSCs using established protocols (61,62) and TH positivity was confirmed by ICC (Supplementary Figure 9A). Mature DAs carrying LRRK2 mutations were treated with the LRRK2 kinase inhibitor MLi-2 for 7 days prior to analysis. Assessment of lysosomal iron revealed a significant increase in both R1441C and Y1699C DA neurons compared to WT controls, which was rescued by MLi-2 treatment (Figure 8A, B). Subsequent lipid peroxidation analyses were focused on R1441C DA neurons, as this mutation showed robust Rab8a phosphorylation (32) and the most pronounced changes across multiple iron-related readouts examined here. R1441C DA neurons exhibited significantly increased lipid peroxidation compared to WT controls (Figure 8C, D). Importantly, the LRRK2 mutation-driven increase in lipid peroxidation was significantly reduced by selective LRRK2 kinase inhibition in DA neurons, with no effect in WT DA neurons (Figure 8C, D). To verify the selectivity of MLi-2 for LRRK2 kinase activity in DA neurons, as previously established in iPSCs (Supplementary Figure 8), LRRK2 KO DA neurons were treated with vehicle or 100 nM MLi-2 for 7 days. MLi-2 had no effect on Liperfluo signal in the LRRK2 KO DA neurons, confirming its specificity for this measure in these neurons (Supplementary Figure 9B, C). Together, these data demonstrate that pathogenic LRRK2 mutations promote lysosomal iron accumulation and lipid peroxidation in human DA neurons, with both phenotypes dependent on a pathogenic increase in LRRK2 kinase activity.

**Figure 8.**
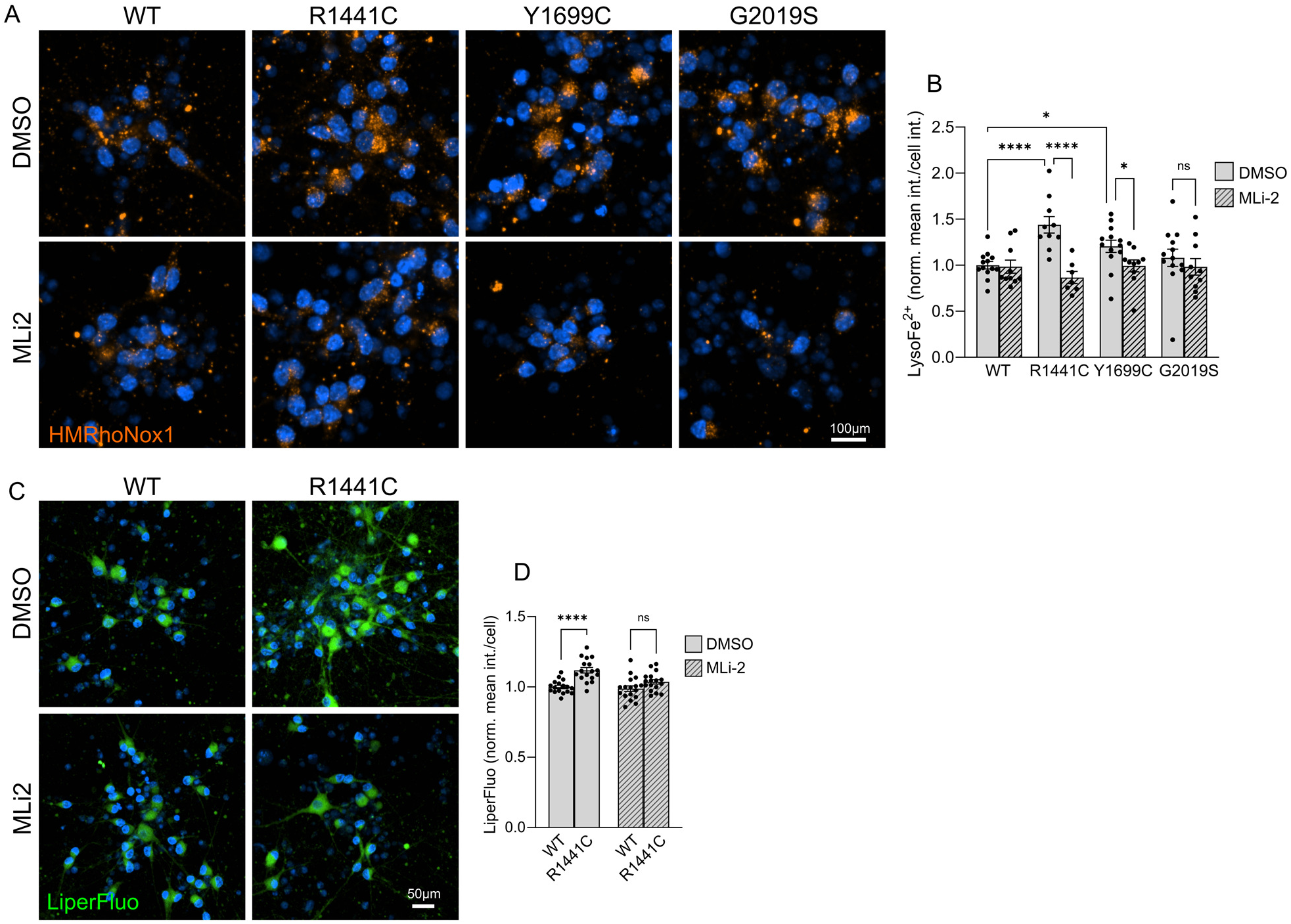
LRRK2 kinase inhibition modulates iron and lipid peroxidation in human dopaminergic neurons. (A, B) Representative images and quantitation of lysosomal iron (HMRhoNox-M) in mature human iPSC-derived DA neurons of the indicated genotypes following treatment with vehicle (DMSO) or MLi-2 (100 nM, 7 days) (two-way ANOVA, Tukey post-hoc; N>10 biological replicates (>800 cells per N), Genotype p=0.1953, Treatment p<0.0001, Interaction p=0.0061). (C, D) Representative images and quantitation of lipid peroxidation (Liperfluo) in WT and heterozygous R1441C DA neurons following treatment with DMSO or MLi-2. (two-way ANOVA, Tukey post-hoc; N=15 biological replicates (>800 cells per N), Genotype p<0.0001, Treatment p=0.0148, Interaction p=0.1211).

## Discussion

The pathological deposition of iron in the PD brain has been widely documented for over three decades, with early histochemical studies identifying iron accumulation in the substantia nigra of post-mortem PD tissue (97,98), and subsequent MRI and spectroscopic studies confirming region-specific iron overload *in vivo* (9,99,100). However, the mechanisms by which genetic risk factors may contribute to intracellular iron mismanagement have remained poorly defined. In this study we show that PD-linked LRRK2 mutations disrupt iron handling in human iPSCs, neurons, and astrocytes, with a striking and convergent increase in lysosomal ferrous iron. This iron accumulation is kinase-dependent and reversible, implicating chronic aberrant LRRK2 signaling in the misregulation of vesicular iron storage. By integrating imaging, biochemical assays, and isogenic models, we report a role for Rab8a in LRRK2-driven iron defects and demonstrate that LRRK2 mutations may enhance iron-driven oxidative stress in human DA neurons. These findings provide a mechanistic link between LRRK2 activity and iron toxicity, offering new insight into the neuronal vulnerabilities that may drive neurodegeneration in PD.

Most pathogenic LRRK2 mutations cluster within the kinase and the Roc-COR tandem domains of the protein, and all are thought to augment the kinase activity of LRRK2 towards a subset of Rab GTPases (34). Rab GTPases control many aspects of intracellular vesicle trafficking by acting as regulatable switches that recruit effector molecules to distinct intracellular membranes (36,101). We and others have shown that phosphorylation of Rab GTPases by mutant LRRK2, including Rab8a and Rab10, interferes with their function and alters endolysosomal dynamics (26,28,32,34,35,102–104). Furthermore, it has been reported that recruitment of LRRK2 to damaged lysosomes, along with Rab8a and Rab10, affects lysosomal integrity (28,32,72). Studies have shown divergent effects of Rab8a and Rab10 in cilliogenesis (105) and we recently reported divergent lysosomal phenotypes between Rab8a KO and Rab10 KO cells (56). Here, we extend this work by demonstrating that loss of Rab8a but not Rab10 partially mimics aspects of iron impairment observed in LRRK2 mutant cells, including upregulation of labile cellular iron and ferritin. Importantly, exogenous Rab8a expression rescued lysosomal iron accumulation in R1441C LRRK2 mutant cells, providing direct evidence that impaired Rab8a activity contributes causally to iron misregulation downstream of pathogenic LRRK2 signaling. Together, these findings define a novel Rab8a-dependent pathway linking LRRK2 kinase activity to intracellular iron handling, potentially through its role in transferrin receptor trafficking and lysosomal function (32,40). The focus on Rab8a is especially interesting given its additional relationship with PINK1 and α-synuclein biology (106–108).

Lysosomes serve as crucial hubs for iron storage and availability (70,71). Physiologic delivery of iron to cells is primarily mediated by transferrin, which binds ferric iron in the extracellular environment and is internalized through clathrin-dependent endocytosis via the transferrin receptor, while non-transferrin bound iron (NTBI) uptake can play a role in iron uptake associated with systemic iron overload or trauma (109). Within endolysosomes, the acidic environment facilitates the release of iron from transferrin and endolysosomal ferrireductases reduce Fe³⁺ to Fe²⁺, allowing iron exit into the cytoplasm through DMT1. This endolysosomal acidification step is critical for iron release and trafficking, and its disruption has been linked to cellular iron deficiency, mitochondrial dysfunction, and inflammation (71,110,111). In fact, inhibition of the lysosomal v-ATPase triggers cellular iron dyshomeostasis, resulting in impaired mitochondrial function and non-apoptotic cell death *in vivo* (110). Here, by using a lysosomal-specific iron probe we found that LRRK2 mutations lead to increased lysosomal ferrous iron content, an effect observed in both terminally differentiated neurons and astrocytes. These data, together with previous reports linking LRRK2 to lysosomal function (28,32,72), and specifically intralumenal pH (26,79), suggest that LRRK2 mutations disrupt normal lysosomal iron handling by altering lysosomal homeostasis. We found that perinuclear lysosomes have a pronounced increase in ferrous iron in LRRK2 mutant astrocytes compared to peripheral lysosomes. Studies have shown that perinuclear lysosomes have lower luminal pH and higher protease activity compared to distal ones (78). Lysosomal positioning correlates with LRRK2 activity towards Rab GTPases with perinuclear lysosomes harboring enhanced kinase-driven Rab signaling (77). We and others have reported dysregulation of lysosomal acidification by mutant LRRK2 in different cellular models (26,31,79,112,113). It is plausible that altered acidification impacts iron mobilization from the lysosome and thus alters overall cellular iron content and availability in the context of LRRK2 mutations.

Iron-induced oxidative stress is a well-documented contributor to neurodegeneration, and our results support a role for LRRK2 mutations in exacerbating this process in DA neurons. Iron-induced cell death, termed ferroptosis, has emerged as a key mechanism contributing to oxidative damage and neuronal loss in PD models (20,86–88). The convergence of aberrant brain iron and lipid peroxidation initiates downstream pathology marked by inflammation, glial dysfunction, demyelination, and ultimately neuronal death (89). Here, we observed increased lipid peroxidation and ROS levels in mutant LRRK2 DA neurons and glutamatergic iNs, which were reversed by iron chelation and LRRK2 inhibition. LRRK2 has been linked to ROS accumulation in different models (50–52), while a recent study reported an effect of LRRK2 mutations in NOX2 activity that in turn has been linked to ferroptosis (51,114,115). Furthermore, studies have linked G2019S LRRK2 with heightened oxidative stress vulnerability in human iPSC-derived DA neurons (116), while more recently, *in vivo* data suggested age-dependent oxidative stress and mitochondrial dysfunction in the SNc of R1441G LRRK2 BAC transgenic mice (117). These data are consistent with mitochondrial abnormalities and dopaminergic alterations reported in G2019S knockin mice (118). It is plausible that dysregulation of the labile and lysosomal iron pools by LRRK2 mutations contributes to the reported effects of LRRK2 on cellular ROS. The lysosomal membrane is exposed to redox-active iron and is an initial target of intracellular oxidant damage, with studies reporting a protective role for intralysosomal iron chelation against ROS damage (119). A recent study identified the lysosomal protein prosaposin as an important regulator of lipid peroxidation and neuronal ROS (92). LRRK2 signaling has been linked to prosaposin function *in vitro* and *in vivo*, highlighting a potential functional interaction that could mediate ferroptosis signaling in this context (120,121). What remains to be established is whether dysregulation of lysosomal Fe^2+^ levels by mutant LRRK2 solely drives cellular ROS damage, or whether aberrant transport of endolysosomal iron to the labile or mitochondrial iron pools also contribute pathologic effects. The finding that deferoxamine attenuated both ROS and lysosomal iron content further strengthens the connection between lysosomal iron mismanagement and oxidative stress in PD-relevant cell types. Ferroptosis inhibitors have shown promise in mitigating neurodegeneration in cell and in *in vivo* preclinical models (20,89,122) Thus, targeting lysosomal LRRK2-mediated iron dysregulation may represent a viable therapeutic strategy.

Cellular iron homeostasis is tightly regulated at the post-transcriptional level by IRPs, which bind to IREs in target mRNAs to control the expression of key iron-handling proteins (80,81,83,84). In our models, ferritin levels were particularly dysregulated in G2019S LRRK2 lines, while IRP1/2 activity showed genotype-specific variation. Notably, cytosolic ferritin levels did not correlate strictly with IRP binding activity, implying that lysosomal dysfunction and impaired ferritin turnover may dominate ferritin regulation in the context of mutant LRRK2 signaling. Interestingly, G2019S showed stronger effects on IRP/ferritin regulation compared to R1441C and Y1699C, but a less convergent effect on iron levels across different cell types. A recent study reported impaired regulation of the ferritinophagy adaptor NCOA4 in G2019 LRRK2 macrophages in a yet-to-be defined kinase-independent manner (123). Previously, we reported elevated levels of ferritin in the brain of G2019S LRRK2 mice, following proinflammatory stimulation (32). It is possible that disruption of ferritinophagy in the context of G2019S LRRK2 underlies altered ferritin levels and mobilization, while ROC-COR domain mutations may impact iron homeostasis via partially divergent mechanisms.

Our data reveal that LRRK2 mutations drive lysosomal iron accumulation in both neurons and astrocytes, supporting a model in which iron dysregulation arises through both cell-autonomous and non-cell-autonomous mechanisms. Astrocytes are essential regulators of brain iron homeostasis, not only by sequestering and buffering excess iron, but also by modulating iron availability at synaptic interfaces through secretion of factors that support neuronal iron uptake (41,42,45–47,124). By shaping the extracellular iron landscape, astrocytes maintain appropriately low synaptic levels and influence neuronal redox balance (45,124). Astrocytes are relevant to LRRK2 biology as they express high levels of LRRK2 (31,125,126), and their cytokine profile is modulated by LRRK2 kinase activity (43,44,48). The increase in lysosomal ferrous iron observed in LRRK2 mutant astrocytes suggests a failure of astrocytic iron buffering capacity, that in an *in vivo* setting would translate into alterations of the synaptic iron microenvironment. In parallel, the accumulation of lysosomal iron in LRRK2 mutant neurons reflects intrinsic defects in iron trafficking and storage, further sensitizing neurons to oxidative injury and ferroptosis. These findings highlight the potential for a dual contribution of LRRK2 to neurodegeneration: through direct, cell-autonomous disruption of neuronal iron handling, and indirectly by impairing astrocyte-mediated regulation of extracellular iron homeostasis.

Based on our current findings from human iPSC-derived models, further work will be needed to confirm whether similar lysosomal iron redistribution occurs *in vivo* in preclinical models where we and others previously reported effects on global iron pathways (32,35,127). Additionally, while iron chelation and LRRK2 inhibition both rescued aspects of the phenotype, a dual therapeutic strategy remains to be tested in preclinical models. Importantly, iron chelation is an effective treatment in Neurodegeneration with Brain Iron Accumulation that often includes parkinsonism symptoms (128–131), but recent clinical trials in PD failed (132) suggesting that targeting precise iron pathways pertaining ferroptosis signaling, in combination with other strategies, in the brain rather than systemic chelation may be considered for future therapeutic efforts for altering disease progression.

## Conclusion

Our findings identify lysosomal iron dysregulation as a key downstream consequence of pathogenic LRRK2 activity in PD. We demonstrate that LRRK2 regulates neuronal and glial iron homeostasis through effects on lysosomal iron trafficking, and that disruption of this pathway leads to redox imbalance and increased ferroptotic vulnerability. The identification of Rab8a, a protein also linked to PINK1 and α-synuclein dependent misregulation, as a critical LRRK2 substrate in this process provides a mechanistic link between LRRK2 signaling and iron biology, offering insight into how genetic and idiopathic PD may converge on a shared pathogenic pathway.

## List of abbreviations

PD: Parkinson’s disease
iPSC: Induced pluripotent stem cell
iNs: Induced neurons
iAs: Induced astrocytes
LRRK2: Leucine-rich repeat kinase 2
ROS: Reactive oxygen species
FAC: Ferric ammonium citrate
DEF: Deferoxamine
TfR1: Transferrin receptor 1
DMT1: Divalent metal transporter 1
Fth1: Ferritin heavy chain
Ftl: Ferritin light chain
IRP: Iron regulatory protein
IRE: Iron-responsive element
ICP-MS: Inductively coupled plasma mass spectrometry
QSM: Quantitative susceptibility mapping
PPMI: Parkinson’s Progression Markers Initiative
ANOVA: Analysis of variance
WT: Wild-type
TH: Tyrosine Hydroxylase

## Declarations

## Ethics approval and consent to participate

Most experiments were performed using isogenic iPSC lines provided by the laboratory of Mark Cookson (National Institutes of Health) that were generated from the commercially available A18945 parental iPSC line (Thermo Fisher Scientific). Written informed consent for research was obtained from all participants in the Parkinson’s Progression Markers Initiative (PPMI), a multi-center study approved by the institutional review boards at each participating site (individual IRB approval numbers are held by the originating institutions). Additional human iPSC lines were obtained from the NINDS Human Cell and Data Repository and were generated under approved institutional review board protocols with documented informed consent. All procedures involving human-derived materials were conducted in accordance with the principles of the Declaration of Helsinki.

## Consent for publication

This manuscript was reviewed by the PPMI Data and Publications Committee and is permitted for publication.

## Availability of data and materials

Materials and additional details can be made available by the corresponding authors upon request.

## Authors’ contributions

Conceptualization: AM, MJL; Methodology: AM, RDB, TBD, AC, NS, AS, NM; Formal analysis and investigation: AM, RDB, TBD, AS, AC, NS, NM; Writing - original draft preparation: AM; Writing - review and editing: AM, MJL, RDB, AC, NS, NM, CDV; Funding acquisition: AM, MJL, CDV; Resources: AM, MJL, CDV; Supervision: MJL, AM. All authors read and approved the final manuscript.

## Acknowledgements

We thank Dr. Jeffrey R. Jones (CTRND, University of Florida) for helpful discussions. We thank the patients and controls who volunteered to the study and the Parkinson’s Progression Markers Initiative (PPMI) for providing access to their samples and data. Data used in the preparation of this article was obtained between 03 November 2023 and 10 November 2023 from the PPMI database (www.ppmi-info.org/access-data-specimens/download-data), RRID:SCR_006431. For up-to-date information on the study, visit www.ppmi-info.org. PPMI, a public-private partnership, is funded by The Michael J. Fox Foundation for Parkinson’s Research and funding partners, including 4D Pharma, AbbVie, AcureX, Allergan, Amathus Therapeutics, Aligning Science Across Parkinson’s, AskBio, Avid Radiopharmaceuticals, BIAL, BioArctic, Biogen, Biohaven, BioLegend, BlueRock Therapeutics, Bristol-Myers Squibb, Calico Labs, Capsida Biotherapeutics, Celgene, Cerevel Therapeutics, Coave Therapeutics, DaCapo Brainscience, Denali, Edmond J. Safra Foundation, Eli Lilly, Gain Therapeutics, GE HealthCare, Genentech, GSK, Golub Capital, Handl Therapeutics, Insitro, Jazz Pharmaceuticals, Johnson & Johnson Innovative Medicine, Lundbeck, Merck, Meso Scale Discovery, Mission Therapeutics, Neurocrine Biosciences, Neuron23, Neuropore, Pfizer, Piramal, Prevail Therapeutics, Roche, Sanofi, Servier, Sun Pharma Advanced Research Company, Takeda, Teva, UCB, Vanqua Bio, Verily, Voyager Therapeutics, The Weston Family Foundation, and Yumanity Therapeutics.

