## Supplementary Figures for "Parkinson’s disease LRRK2 mutations dysregulate iron homeostasis and promote oxidative stress and ferroptosis in human neurons and astrocytes"

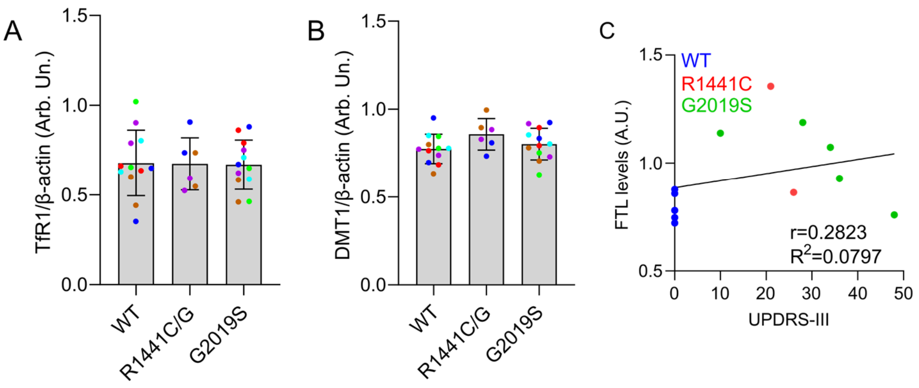


**Supplementary Figure 1.** (A, B) Western blot and quantitation of levels of iron-related factors in WT, heterozygous R1441C/G, and G2019S iPSCs (N=2 biological replicates per line; 6 WT, 2 R1441G, 1 R1441C and 6 G2019S lines, one-way ANOVA, Dunnett's post-hoc). (C) Scatter plot showing no significant correlation between FTL levels and clinical severity of patient donors (UPDRS-III).

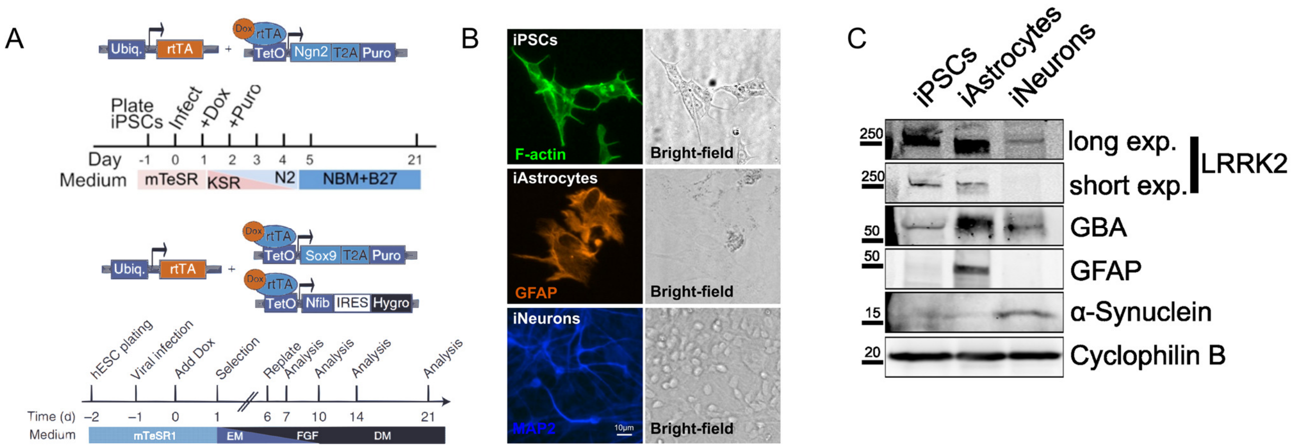


**Supplementary Figure 2**. iN and iA differentiation protocols and validation of neuronal and astrocytic markers. (A) iPSCs were differentiated into iNs by forced expression of NGN2, and into iAs by forced expression of SOX9 and NFIB, according to published protocols. (B, C) Expression of MAP2, GFAP and α-synuclein was validated by ICC and immunoblotting.

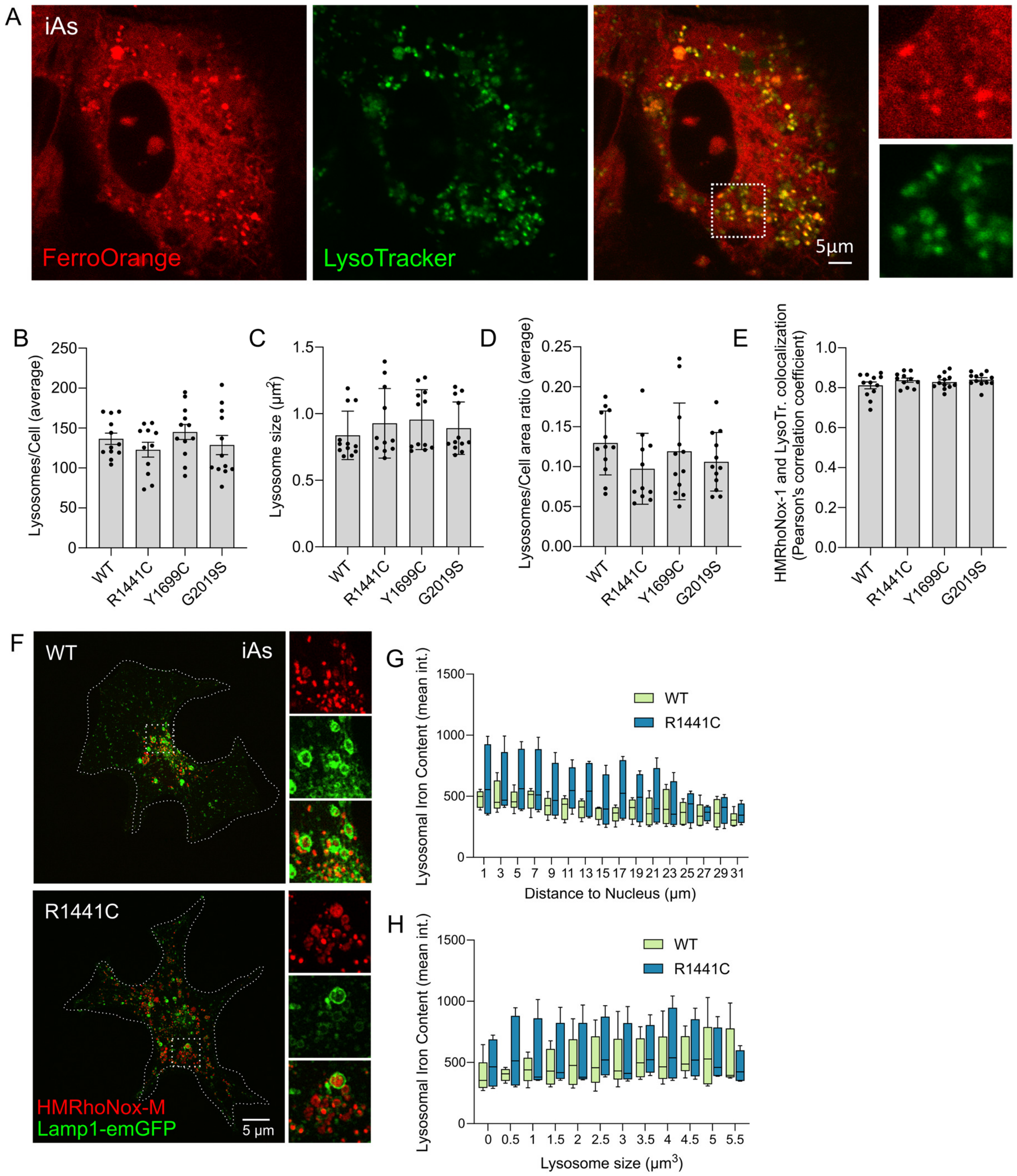


**Supplementary Figure 3. Characterization of HMRhoNox-M dye and lysosomes in iAs.** (A) iAs were stained with the FerroOrange iron probe and LysoTracker dye and imaged by confocal microscopy. (B-D) Quantitation of lysosomal number and size across mutant LRRK2 iAs and isogenic controls. (E) Quantitation of colocalization between HMRhoNox-M and LysoTracker across iAs carrying LRRK2 mutations and isogenic controls. (F) Super-resolution imaging of lysosomal iron and Lamp1-emGFP in WT and isogenic R1441C iAs. Z-stack confocal images were 3D reconstructed in Imaris (Oxford Instruments) and the frequency distribution of lysosomal iron content versus proximity to the nucleus (G) as well as lysosome size (H), were plotted.

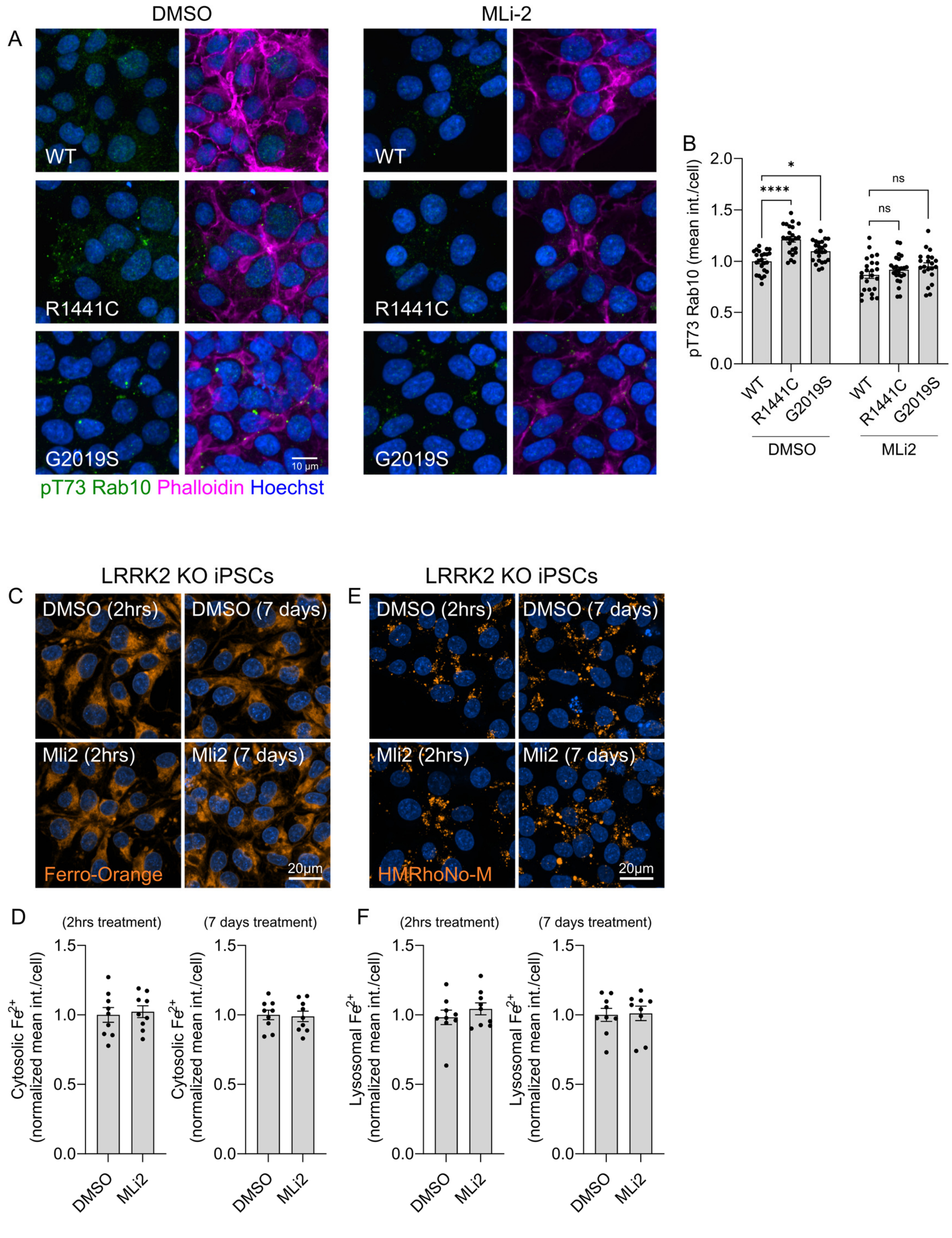


**Supplementary Figure 4. Validation of LRRK2 kinase inhibition and specificity of MLi-2 effects on iron homeostasis.** (A) Representative images and quantitation of phosphorylated Rab10 at T73 in isogenic WT, R1441C, and G2019S iPSCs treated with vehicle (DMSO) or MLi-2 (100 nM) for 2 hours. Representative images and quantitation of cytosolic (FerroOrange; C, D) and lysosomal iron (HMRhoNox-M; E, F) in LRRK2 KO iPSCs treated with DMSO or MLi-2 for either 2 hours or 7 days. (pT73: one-way ANOVA with Dunnett's post hoc test; FerroOrange and HMRhoNox-M: student's t-test).

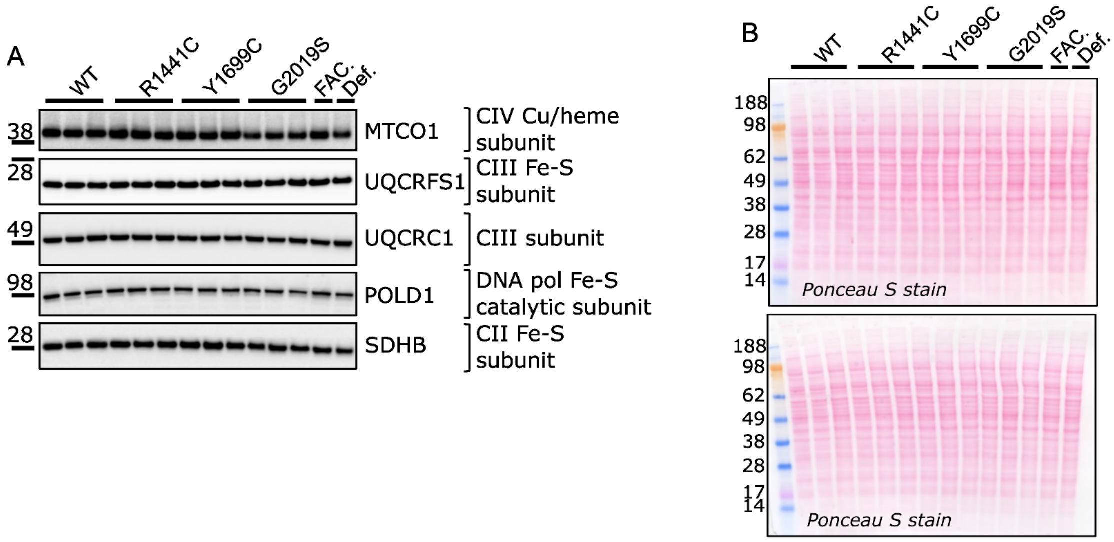


**Supplementary Figure 5. Expression of Fe-S cluster containing proteins in iPSCs.** (A) Isogenic iPSCs carrying LRRK2 mutations were analyzed for expression of Fe-S containing factors by immunoblot. (B) Equal loading was further validated by ponceau staining.

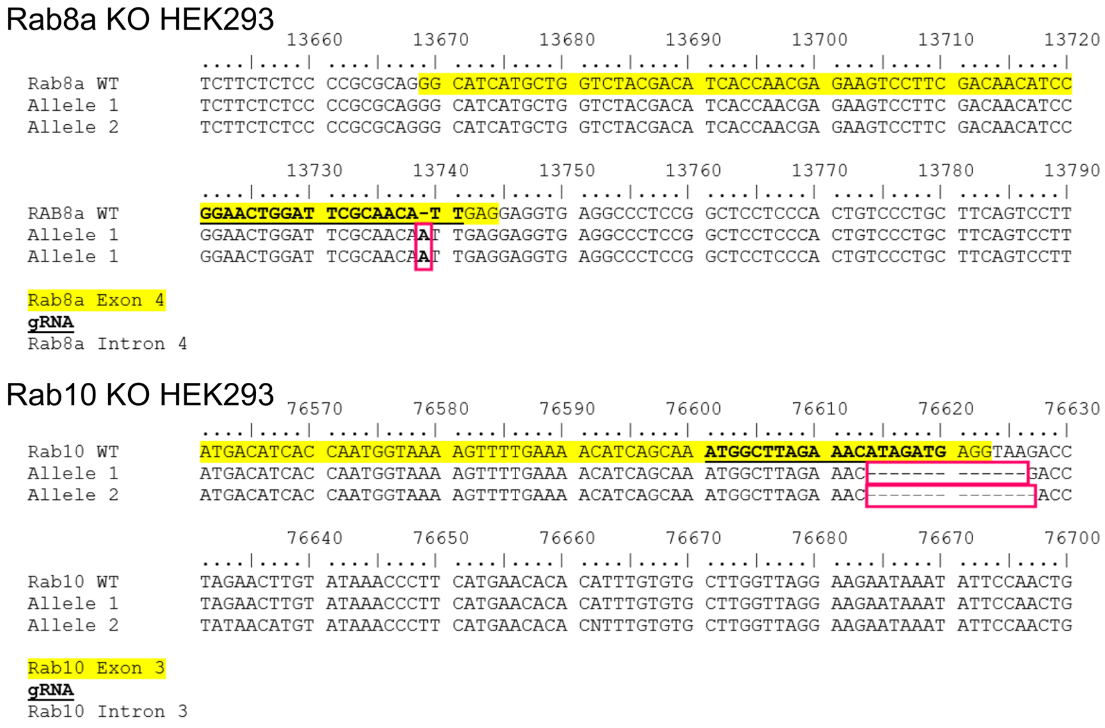


**Supplementary Figure 6. Validation of *RAB8a* KO and *RAB10* KO HEK293 clones by Sanger sequencing.** HEK293 Clones of each *RAB* GTPase edited line were validated by Sanger sequencing and used for further experimentation.

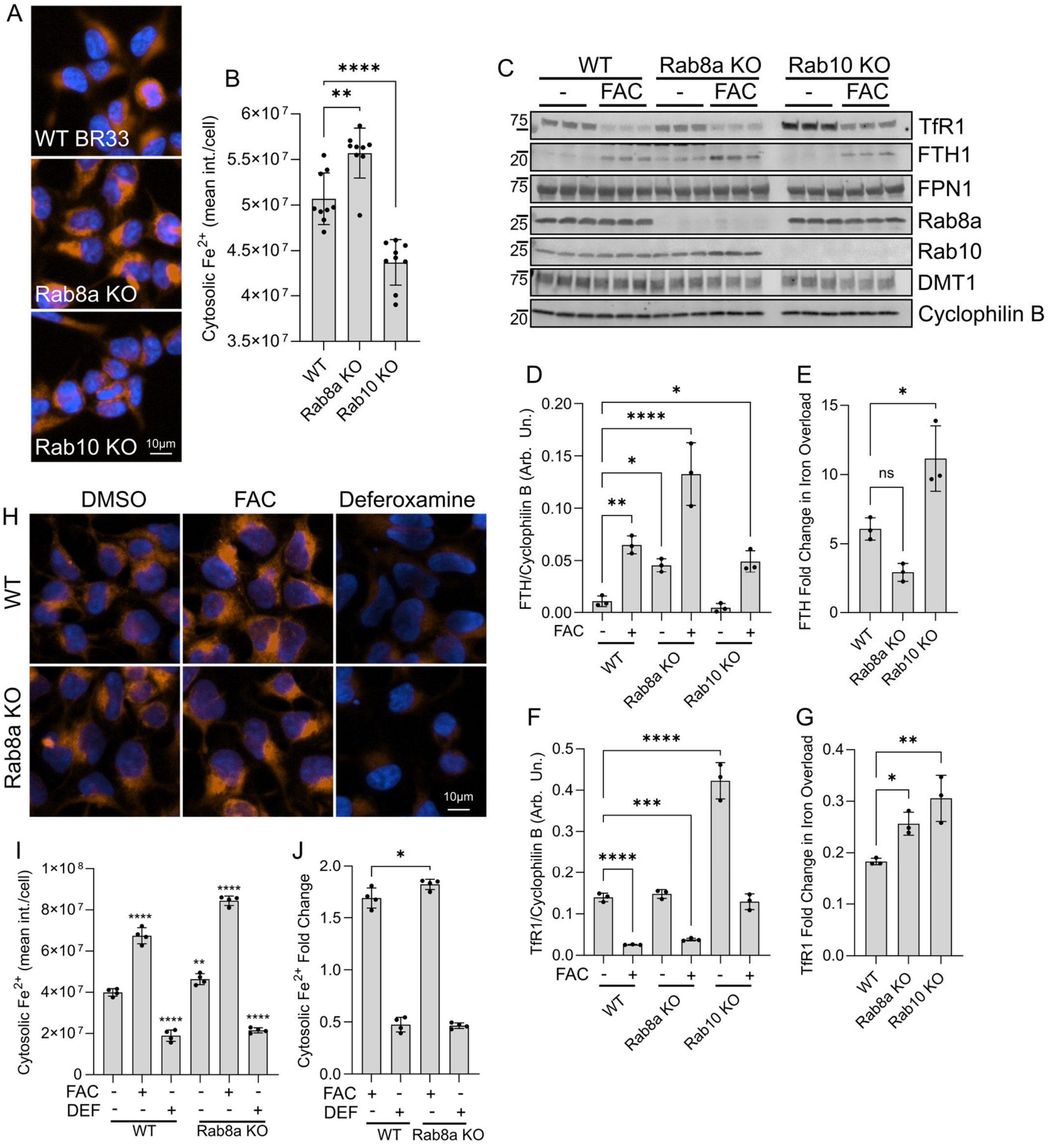


**Supplementary Figure 7. Rab8a deficiency impairs ferritin heavy chain regulation.** (A, B) Imaging of cytosolic free iron and quantitation of iron levels in WT, Rab8a KO and Rab10 KO iPSCs by high-content imaging. (one-way ANOVA, Dunnett's post-hoc, N=6 biological replicates, *p<0.05, **p<0.01, ****p<0.0001). (C-G) Western blot analysis of iron-related proteins in Rab8a KO and Rab10 KO HEK293 cells in basal and iron overload (FAC) conditions (two-way ANOVA, Tukey's post-hoc; N=3 biological replicates; Fth1: Genotype ****p=0.0001, Treatment ****p<0.0001, Interaction p=0.0508; TfR1: Genotype ****p=0.0001, Treatment ****p<0.0001, Interaction ****p<0.0001). (H-J) Imaging and quantitation of free iron in Rab8a KO HEK293 cells under iron overload (FAC) and iron chelation (DEF) conditions. (one-way ANOVA, Dunnett's post-hoc, N=4 biological replicates, *p<0.05, **p<0.01, ***p<0.001).

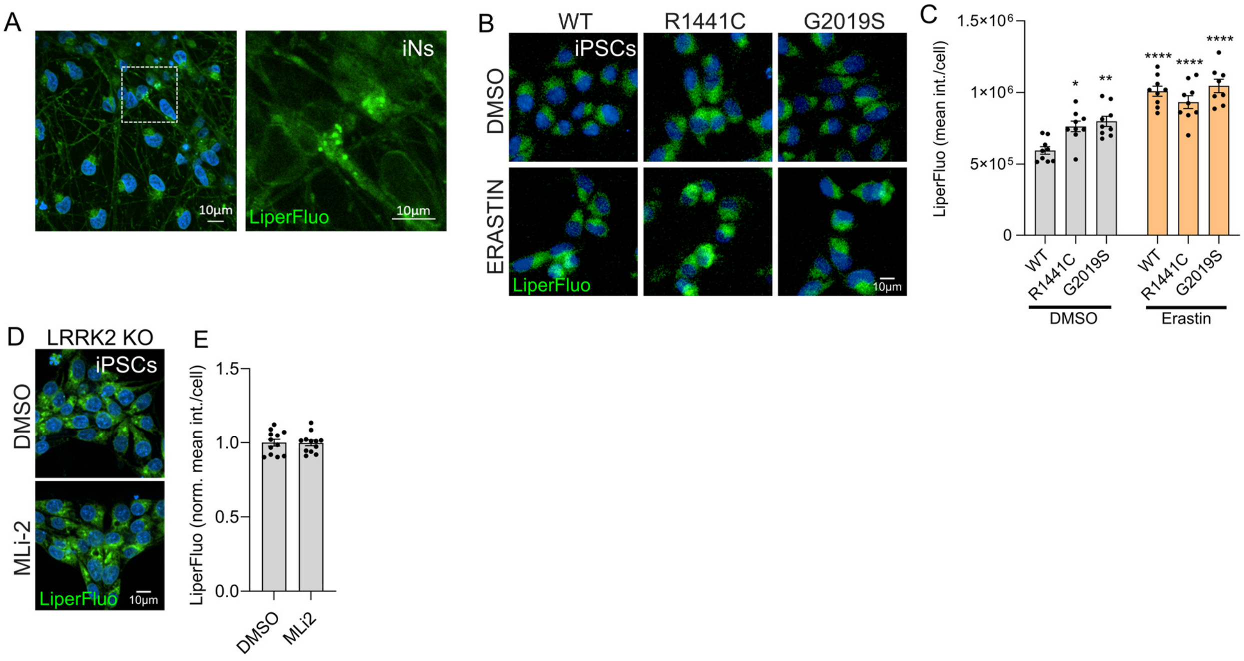


**Supplementary Figure 8. LRRK2 mutations increase basal lipid peroxidation.** (A) Representative confocal image of iNs stained with Liperfluo, illustrating punctate lipid peroxidation signal. (B) Isogenic iPSCs carrying LRRK2 mutations stained with Liperfluo to detect lipid peroxidation following treatment with Erastin or vehicle. (C) Quantitation of Liperfluo intensity by high-content imaging (two-way ANOVA, Tukey's post-hoc; N=9 biological replicates (>800 cells per N), Genotype *p=0.0105, Treatment ****p<0.0001, Interaction **p=0.0072). (D, E) Representative images and quantification of Liperfluo signal in LRRK2 KO iPSCs treated with DMSO or MLi-2 (100 nM, 7 days). MLi-2 did not alter lipid peroxidation in LRRK2 KO cells, confirming specificity of LRRK2-dependent effects observed in mutant neurons.

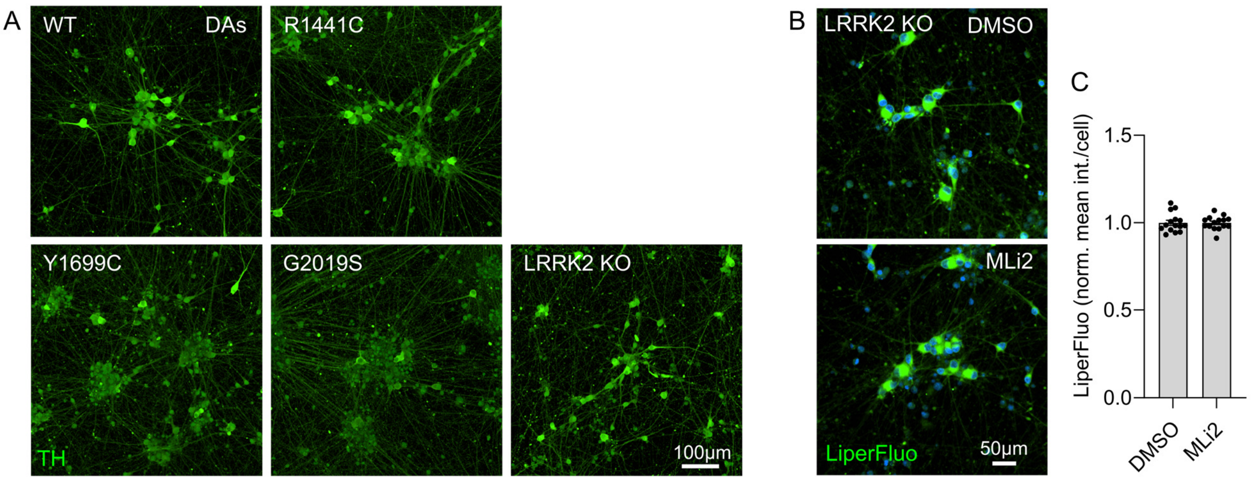


**Supplementary Figure 9. DA neuron differentiation and specificity of MLi-2 effects in DA neurons.** (A) Representative images of TH immunostaining in human iPSC-derived DA neuron cultures of WT and isogenic LRRK2 mutant lines. (B, C) Representative images and quantitation of Liperfluo staining in LRRK2 KO DA neurons treated with DMSO vehicle or MLi-2 (100 nM, 7 days).

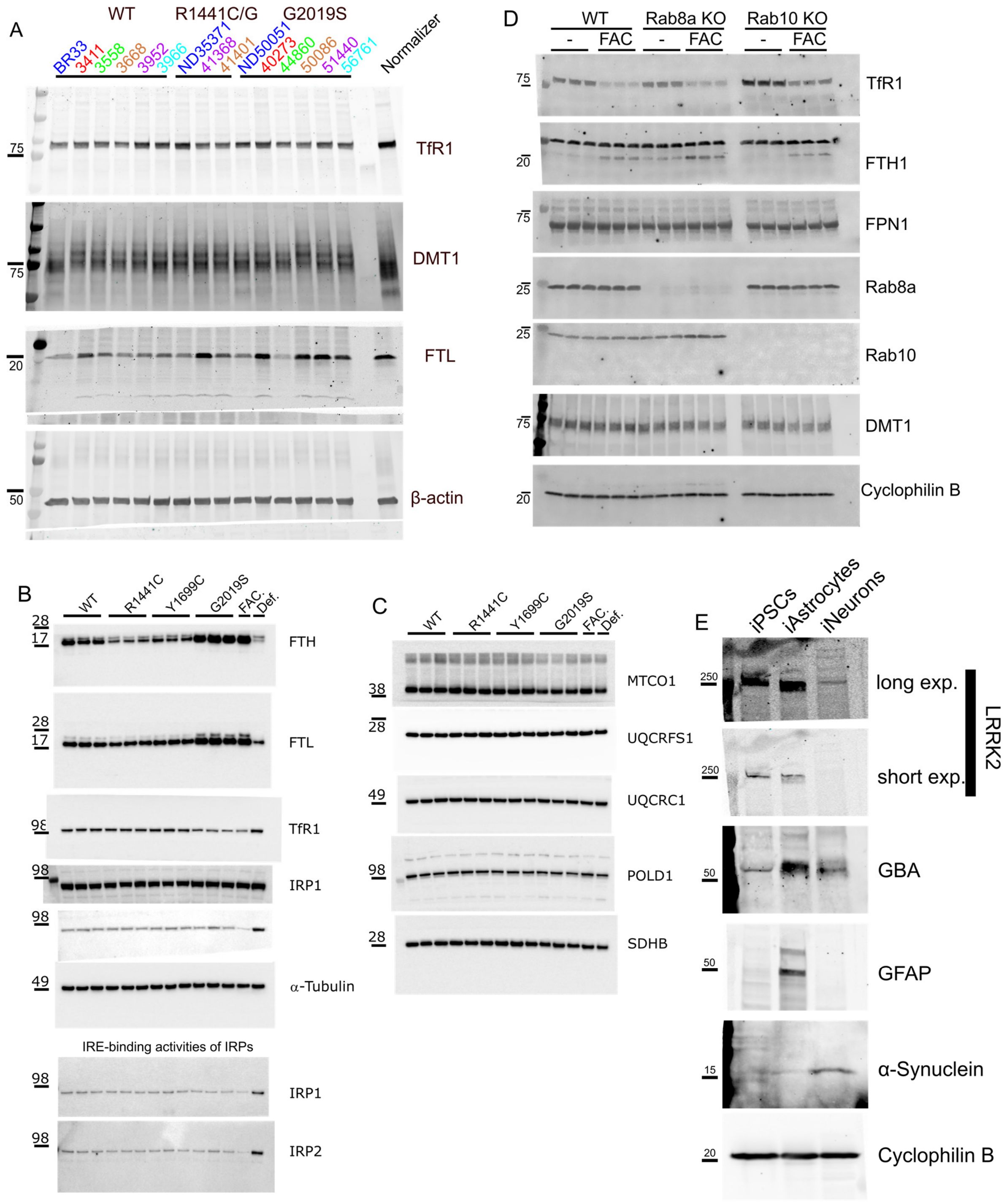


**Supplementary Figure 10. Uncropped blots.** Full blots are presented from: (A) Figure 1F, (B) Figure 3A, (C) Suppl. Figure 2C, (D) Suppl. Figure 3A.
